# Cholinergic modulation of reinforcement learning and prefrontal value computations under uncertainty

**DOI:** 10.1101/2024.09.20.614105

**Authors:** Hannah Kurtenbach, Monja Isabel Froböse, Eduard Ort, Bahne Bahners, Markus Butz, Alfons Schnitzler, Jan Hirschmann, Gerhard Jocham

## Abstract

The neuromodulator acetylcholine has been suggested to govern learning under uncertainty. Here, we investigated the role of muscarinic acetylcholine receptors in reward-guided learning and decision making under different degrees of uncertainty. We administered the muscarinic antagonist biperiden (4 mg) to healthy male participants (n = 43) in a within-subjects, placebo-controlled design. Participants performed a gambling and a learning task involving different levels of uncertainty, while magnetoencephalography (MEG) was recorded. We show that biperiden did not affect decision making in the gambling task, where no learning was required. However, in the learning task, where option values were associated with uncertainty, biperiden reduced the sensitivity to probabilities, particularly when choice-outcome-contingencies switched frequently. Reinforcement learning models revealed that the change in behaviour was caused by noisier estimates of probabilities resulting from increased learning rates for rewarded choices under biperiden. These behavioural findings were paralleled by elimination of the lateral prefrontal representation of learnt reward probability in high-beta (20 - 30 Hz) power under biperiden. Together, these findings suggest that muscarinic acetylcholine transmission controls learning in highly uncertain contexts, when the demand for carefully calibrated adjustments is highest.

## Introduction

Decision making usually involves some degree of uncertainty. Often, information relevant for a choice is not known and needs to be inferred from experience. In addition, choice-outcome contingencies may be probabilistic and even change over time. When option attributes are not explicitly presented and have to be learnt from trial and error, choice behaviour has been formally described by reinforcement learning algorithms (Daw et al., 2011; Kurtenbach et al., 2022; Lee et al., 2012; Sutton & Barto, 2014). Core to these algorithms is the updating of value estimates using the prediction error, the discrepancy between obtained and expected outcomes. This prediction error is scaled by a learning rate parameter that determines the degree to which the error is used to update value estimates. In real life, the link between choices and outcomes is often fraught with uncertainty, which requires agents to estimate this uncertainty for adaptive decision making (Behrens et al., 2007; Lee et al., 2012). It has been shown that participants adjust learning rates to the statistics of the environment (Iglesias et al., 2021; Jocham et al., 2009; Soltani & Izquierdo, 2019). In particular, learning rates have been shown to increase in volatile environments when contingencies change frequently (Behrens et al., 2007; Browning et al., 2015).

Acetylcholine is a neurotransmitter that has been suggested to play an important role in learning under uncertainty. A long tradition of research, in particular using pharmacological and lesion approaches, has firmly established a role for basal forebrain cholinergic neurons in learning and attention (reviewed in e.g. Everitt and Robbins, 1997; Hasselmo and Sarter, 2011). Beyond this, theoretical work proposes that acetylcholine controls the learning rate (Doya, 2002) and governs decision making in environments with known uncertainty (Avery et al., 2012; Yu & Dayan, 2005). In addition to this presumed role in uncertainty-dependent learning, acetylcholine modulates neural circuit dynamics supporting reward-guided choice, irrespective of learning: Activation of cholinergic receptors, especially the muscarinic M1 subtype, enhances the function of NMDA glutamate receptors and activates GABAergic inhibitory circuits (Bessie Aramakis et al., 1997; Kuchibhotla et al., 2017; Marino et al., 1998; Obermayer et al., 2017; Zwart et al., 2018). These two neurotransmitter systems are the key components in recurrent cortical circuit models of decision making, where competition between options is governed by slow excitation at NDMA receptors and GABAergic feedback inhibition (Wang, 2002; Wong & Wang, 2006). This suggests that acetylcholine may affect decision-making computations beyond its presumed role in learning.

At the circuit level, the computations underlying decision making are reflected in neural oscillations in different frequency bands. Evidence accumulation has been related to prefrontal beta and gamma oscillations (Polanía et al., 2014), whereas outcome evaluation and reward processing are reflected in theta- and high-beta power (Christie & Tata, 2009; HajiHosseini & Holroyd, 2015; Marco-Pallares et al., 2008; Van De Vijver et al., 2011). In particular, high-beta band activity (20 – 30 Hz) has been reported to increase following reward feedback and to reflect the evaluation of positive outcomes (Marco-Pallares et al., 2008; Marco-Pallarés et al., 2015). Moreover, beta-band oscillations have been proposed to signal the maintenance of internal models and top-down control following feedback (Engel & Fries, 2010; Tan et al., 2014). Acetylcholine, in turn, has been shown to modulate neural oscillations, among others, in the beta band (Bauer et al., 2012), possibly via modulation of GABAergic interneurons that shape beta oscillations (Chen et al., 2015; Kuchibhotla et al., 2017; Veit et al., 2017). Together, this suggest that acetylcholine may profoundly affect the local circuit and network computations underlying learning and decision making.

The current study aimed to investigate the behavioural and neural effects of muscarinic acetylcholine receptor antagonism in both decision-making computations and learning, using paradigms probing reward-guided decision making under varying degrees of uncertainty. To this end, participants performed two reward-guided choice tasks under the influence of either placebo or the muscarinic receptor antagonist biperiden (4 mg) while cortical activity was recorded using magnetoencephalography (MEG). The gambling task involved explicit reward probability information, whereas in the learning task reward probabilities had to be inferred from experience during phases with either stable or changing (i.e. volatile) choice-outcome contingencies. We hypothesized that muscarinic antagonism would impair decision making and learning across both tasks. Notably, we found that biperiden had no effect on choice behaviour in the gambling task, but it affected decision making in the learning task, in particular when reward-contingencies frequently reversed. Biperiden increased the learning rate in the volatile phase of the learning task in a maladaptive manner, as evident from noisier estimates of the underlying reward probability. These behavioural findings were accompanied by a disruption of prediction error representations in lateral prefrontal cortex high-beta power under biperiden.

## Results

We administered the M1-preferring muscarinic receptor antagonist biperiden (4 mg) to healthy male participants (n = 43) and acquired MEG in a within-subjects, placebo-controlled design (Fig. 1a). Participants performed two reward-guided choice tasks with the goal to maximize their reward by selecting one option from a pair of gambles each associated with a reward probability and reward magnitude. In the gambling task (Fig. 1b), both attributes were explicitly presented to participants. The reward magnitude was provided by the height of rectangular bars and the reward probability was presented numerically underneath each bar. Whether or not the reward was paid out depended on the explicit reward probability. In the learning task (Fig. 1b), reward magnitudes were again explicitly expressed by the height of the rectangular bars, but here, reward probability of the two options (indicated by two colours) had to be learnt from experience. One colour had a reward probability of 0.7, the other of 0.3. To evoke different levels of uncertainty, the learning task featured a stable and a volatile phase. In the stable phase, the mapping of reward probabilities to the two colours was fixed, whereas in the volatile phase, the contingencies between reward probability and colour reversed several times over the course of the experiment (Fig. 1c).

**Figure 1.**
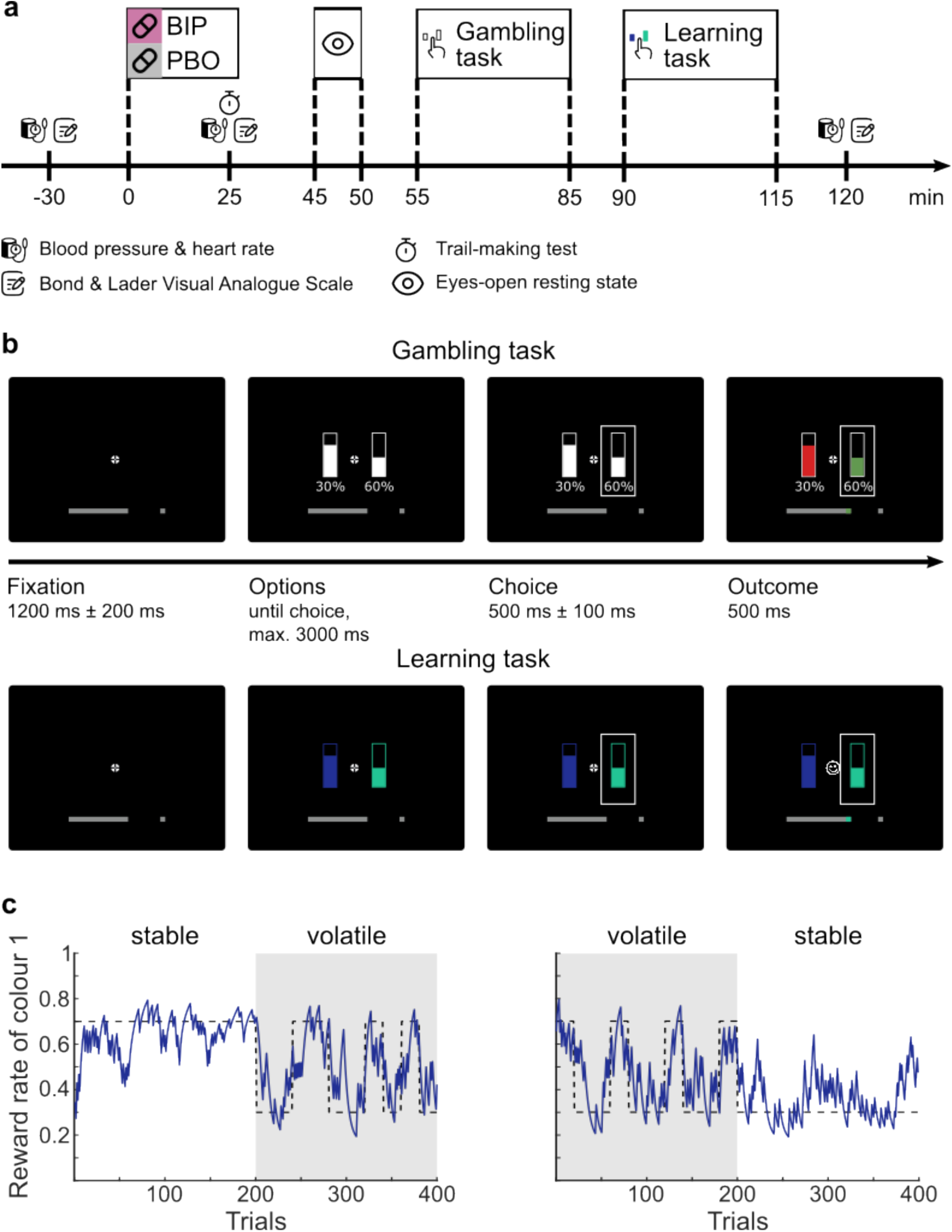
Schematic of study procedure and experimental tasks. **a** Study procedure. Participants received either the biperiden (BIP) or placebo (PBO) pill. 45 minutes after drug, a resting state MEG scan was acquired. The gambling task was scheduled to start 55 minutes (55.9 min ± 1.7 min) and the learning task approximately 90 minutes (91.3 min ± 3.6 min) after drug intake. **b** Example trial of the gambling and the learning task. In both tasks reward magnitudes were indicated by the height of the bar. In the gambling task, reward probabilities were indicated by the numeric percentage below each bar. Outcomes were presented as fill colour in green or red to indicate win or no win, respectively. In the learning task, reward probabilities were indicated by the colour of the bars and needed to be learnt during the task. Outcome was presented as smiley when the winning option was chosen and as frowny otherwise. **c** Example time course of reward contingencies during the learning task. Reward contingencies either remained stable for 200 trials or switched every 20 to 40 trials (volatile phase). Dashed black lines represent true underlying probabilities, solid blue lines represent probability estimates derived from a statistically optimal Bayesian learner.

### Biperiden modulates sensitivity to learnt values

To quantify how task manipulations and cholinergic intervention affected choices in both tasks, we used logistic mixed-effects regressions (see supplementary Fig. S1 for raw behaviour). For these analyses, reward probabilities and magnitudes were entered as regressors in addition to the expected values, a measure of the multiplicative attribute integration, above and beyond the main effects of probability and magnitude.

For the learning task, reward probabilities were estimated using a Bayesian optimal learner (Behrens et al., 2007), because participants could not know the true reward probabilities at the outset of the task. Thus, these probability estimates reflect what the participants could optimally know on a given trial. In line with previous work (Dias Maile et al., 2024; Farashahi et al., 2019; Jocham et al., 2012, 2014), all task parameters had a significant main effect on choice in the gambling task (probability: *β* = 2.90, *SE* = 0.03, *z* = 95.78, *p* < .001; magnitude: *β* = 1.58, *SE* = 0.02, *z* = 73.66, *p* < .001; expected value: *β* = 0.50, *SE* = 0.02, *z* = 30.05, *p* < .001, Fig. 2a, left). In the learning task, choice behaviour was significantly driven by probability and magnitude (probability: *β* = 1.27, *SE* = 0.02, *z* = 80.28, *p* < .001; magnitude: *β* = 0.75, *SE* = 0.01, *z* = 51.45, *p* < .001; Fig. 2a, right, see supplementary Tables S1 and S2 for all results). Surprisingly, expected value had a negative effect, although very small (*β* = -0.04, *SE* = 0.02, *z* = -2.30, *p* = .021).

**Figure 2.**
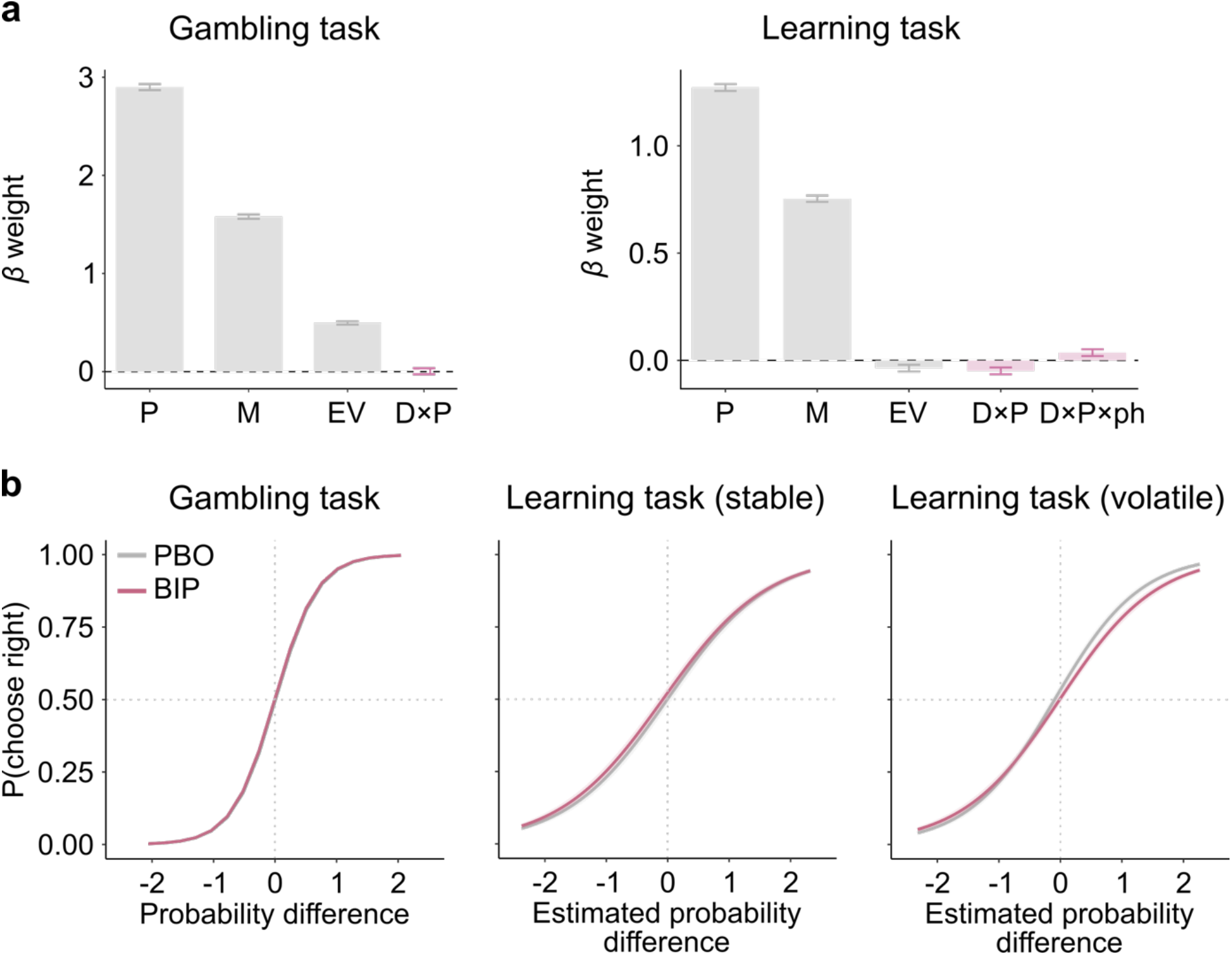
Results from logistic mixed-effects regressions. **a** Task (probability P, magnitude M, and expected value EV) and drug effects for the gambling task (left) and the learning task (right). The interaction between drug (D) and probability (P) is depicted for both tasks, the interaction between drug, probability and phase (ph) is shown for the learning task. **b** Interaction between reward probability and drug. Probability for a right-side choice as a function of differences in reward probability (right vs. left option) for the gambling task, and for the stable (middle) and the volatile phase (right) of the learning task. Post-hoc tests following up a significant interaction indicate that, in the volatile phase only, sensitivity to reward probabilities was significantly reduced under biperiden (BIP, pink) compared to placebo (PBO, grey). Solid lines represent mean, shaded areas 95 % CI.

Contrary to our expectation, biperiden diminished the effect of probability on choice only in the learning task, while leaving choices in the gambling task unaffected. Under biperiden, participants relied less on probability (*β* = -0.05, *SE* = 0.02, *z* = -3.07, *p* = .002). This drug effect was dependent on phase (drug x probability x phase: *β* = 0.04, *SE* = 0.02, *z* = 2.29, *p* = .022). Post-hoc tests indicated that there was no significant effect of drug on probability weighting in the stable phase, whereas in the volatile phase, the use of probability information was significantly reduced under biperiden (*β* = -0.08, *SE* = 0.03, *z* = -3.36, *p* < .001; Fig. 2b). Control analyses revealed that, even though biperiden significantly reduced heart rate and subjective reports of alertness, calmness, and contentedness on the Bond and Lader Visual Analogue Scale (BL VAS; Bond & Lader, 1974; see supplementary Tables S3-S8, Fig. S2-S4), the biperiden-induced reduction in sensitivity to reward probability in the volatile phase was independent of these effects (see supplementary Table S9). Similarly, the effect was independent of the order of stable versus volatile phases (which varied between participants), independent of session days, and independent of the order of drug (see supplementary Table S9).

In sum, logistic mixed-effect regressions revealed that cholinergic modulation of choice behaviour occurred only in the volatile phase of the learning task. We observed that, under biperiden, participants made less use of reward probability information in the volatile phase.

### Biperiden increases learning rate in highly uncertain environments

Logistic mixed-effects regressions revealed that biperiden changed the influence of the implicit reward probability on choice behaviour in the volatile phase of the learning task. However, this probability estimate is derived from a Bayesian optimal learner and thus reflects what participants could ideally know. It remains unclear whether participants tracked probabilities in a statistically optimal manner. Thus, rather than reflecting a diminished use of probability information under biperiden, our result could also indicate a failure to optimally track probabilities. These two mechanisms are not mutually exclusive, they might jointly contribute to our behavioural findings. To adjudicate between these possible mechanisms, we fitted Bayesian hierarchical models that captured (i) the use of learnt reward probability information via relative attribute weighting (magnitudes vs. probabilities) and (ii) probability tracking using reinforcement learning, where learning rates determine the degree to which prediction errors are used to update probability estimates on each trial. Specifically, subjective values for each option are constructed by adding probabilities and magnitudes for one option, after scaling both attributes by a parameter *ω*_*P*_ which indicates the degree to which participants rely more on probabilities relative to magnitudes, or vice versa (methods, equation 3). Reward probabilities were estimated using Q-learning with a delta update rule with learning rates *λ*_*r*_ and *λ*_*u*_, for rewarded and unrewarded choices. Subjective values were transformed into model choice probabilities for the two options using softmax action selection with an inverse temperature parameter. Informed by the regression results reported above, we focus here on drug-induced effects in the volatile phase of the learning task. In the supplementary materials, we also report both drug-independent adaptations to uncertainty (Fig. S5, for a detailed comparison of choice strategies across tasks), and biperiden-induced shifts for the gambling task (Fig. S6, Table S10) and the stable phase of the learning task (Fig. S7, Table S11).

To test for cholinergic effects on these model parameters, we extended the Bayesian hierarchical models with a biperiden-specific shift in each fitted model parameter: A shift of (i) the relative attribute weighting towards magnitudes would indicate that biperiden may have diminished the impact of the learnt (uncertainty-laden) attribute on choice, whereas an effect on (ii) the learning rate (rather than on choice) would indicate that, under biperiden, the accuracy of participants’ probability estimate is altered in the first place. The models revealed that there was a credible biperiden-induced increase of the learning rate for rewarded choices *s*_*λr*_ (HDI_mdn_ = 0.09, HDI_.95_ = [0.01, 0.16], Fig. 3a), but not for the learning rate for unrewarded choices *s*_*λu*_, attribute weighting *s*_*ωP*_, or inverse temperature s _*ζ*_ (see table 1, Fig. 3b-d). Increased learning rates are generally considered to be more adaptive in volatile environments (Behrens et al., 2007; Browning et al., 2015). To capture what participants could ideally know about the underlying reward probabilities, we used a Bayesian optimal learner (Behrens et al., 2007) and explored to what extent estimated probabilities using participants’ fitted learning rates differed from this optimal learner’s estimates. We then computed the mean squared difference between these two estimates and found it to be increased under biperiden compared to placebo (Δ(MSE_BIP_ – MSE_PLA_) = 9.25×10^-3^ ± 0.21×10^-3^, *t*(42) = 6.61, *p* = 0.009, Cohen’s *d* = 1.01, Fig. 3e). This indicates that a further increase of the already rather high learning rates (median(*λ*_*r*_) = 0.58) is maladaptive, as it caused noisier value estimates of implicit information.

**Table 1.** Group-level parameter estimates of the Bayesian hierarchical model for the volatile phase of the learning task. Median (Mdn), standard deviation (SD), and lower and upper bounds of the 95 %-HDI interval are presented. The model included a learning rate of rewarded choices *λ* _*r*_, a learning rate of unrewarded choices *λ* _*u*_, attribute weighting *ω* _*P*_, a softmax inverse temperature *σ*, and the corresponding biperiden-specific shifts on these parameters *s*_*λr*_, *s*_*λu*_, *s*_*ωP*_, and *s*_*ζ*_ .

| Parameter | Mdn | SD | 2.5 % | 97.5 % |
| --- | --- | --- | --- | --- |
| $\lambda_r$ | 0.58 | 0.03 | 0.52 | 0.64 |
| $\lambda_u$ | 0.26 | 0.02 | 0.22 | 0.31 |
| $\omega_P$ | 0.59 | 0.04 | 0.51 | 0.67 |
| $\zeta$ | 5.60 | 0.40 | 4.83 | 6.40 |
| $s_{\lambda_r}$ | 0.09 | 0.04 | 0.01 | 0.16 |
| $s_{\lambda_u}$ | 0.02 | 0.01 | 0.00 | 0.04 |
| $s_{\omega_P}$ | -0.03 | 0.02 | -0.08 | 0.01 |
| $s_{\zeta}$ | -0.37 | 0.23 | -0.81 | 0.10 |

**Figure 3.**
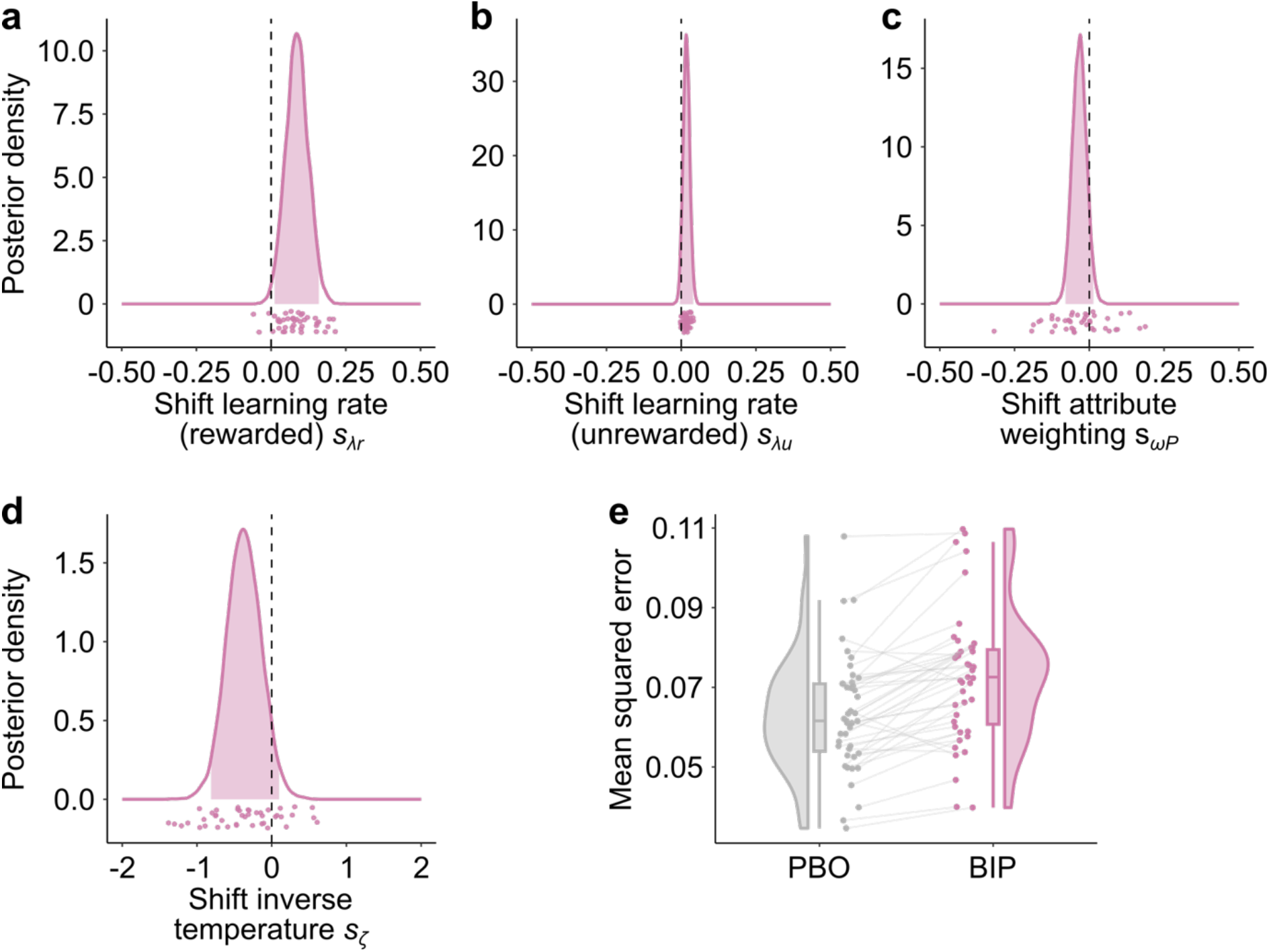
Biperiden-induced shifts of model parameters in the volatile phase of the learning task, derived from the Bayesian hierarchical model. Density of posterior distributions of the biperiden-specific shift in **a, b** learning rate in rewarded *s*_*λr*_ and unrewarded trials *s*_*λu*_, **c** attribute weighting *s* _*ω P*_, and **d** inverse temperature *s* _*ζ*_ . Positive shifts represent an increase under biperiden relative to placebo. The learning rate for rewarded choices is credibly increased under biperiden. Shaded areas represent the 95 %-HDI of the posterior distribution, points are single-subject means. **e** Mean squared error (MSE) of the deviation of participants’ estimated reward probabilities from the estimate obtained from a Bayesian optimal learner, plotted for the placebo session (PBO; grey) and biperiden session (pink). Points reflect individual participants’ fit. Under biperiden, the MSE is significantly higher, reflecting impaired estimation of the learnt attribute.

Taken together, these results indicate that the diminished impact of probability on choice under biperiden observed in the logistic regression does not result from participants using this information less to guide their choices. Instead, acetylcholine appears to be involved in tracking reward probabilities via appropriately setting the learning rate, particularly under conditions of high uncertainty.

### Biperiden modulates prefrontal high-beta-band dynamics

Our behavioural results indicate that biperiden impaired use of outcomes to update probability estimates for future choices in the volatile phase of the learning task. In a first step, we identified a drug-independent region of interest (ROI) showing responses to outcomes. To this end, we calculated the average high-beta-band power for rewarded versus unrewarded choices, averaged over both sessions of each participant, for the interval of 0 – 1000 ms post outcome (Fig. 4a). We focused on oscillations in the high-beta band (20 – 30 Hz) due to their established role in outcome processing (e.g. Andreou et al., 2017; Cohen et al., 2007; HajiHosseini & Holroyd, 2015; Kawasaki & Yamaguchi, 2013; Marco-Pallares et al., 2008), but for exploratory purposes, we also visualized reward responses in the theta, alpha, low-beta, and low-gamma band; see supplementary Fig. S8). We observed a pronounced reward response in the right dorsolateral prefrontal cortex (superior and middle frontal gyrus, Fig. 4a) and performed all of our subsequent analyses on this ROI (Fig. 4b-c). Next, to quantify how biperiden modulates reward responses as a function of task parameters, we extracted trial-wise high-beta-band power from this ROI, averaged over the time window from 0 to 1000 ms after feedback onset. We then entered high-beta power as dependent variable into mixed-effects linear regressions. Our key predictors were drug (biperiden vs. placebo) and the prediction error, the term that, scaled by the learning rate, is used to update probability estimates. Since prediction errors are strongly correlated with outcomes, we directly decomposed the prediction error by entering separate regressors for the outcome (reward vs. non-reward), the subjective probability estimates and their interaction with drug into the model. To account for motor-related beta-band activity, we also included motor response (left vs. right) on the current and previous trial in the model.

**Figure 4.**
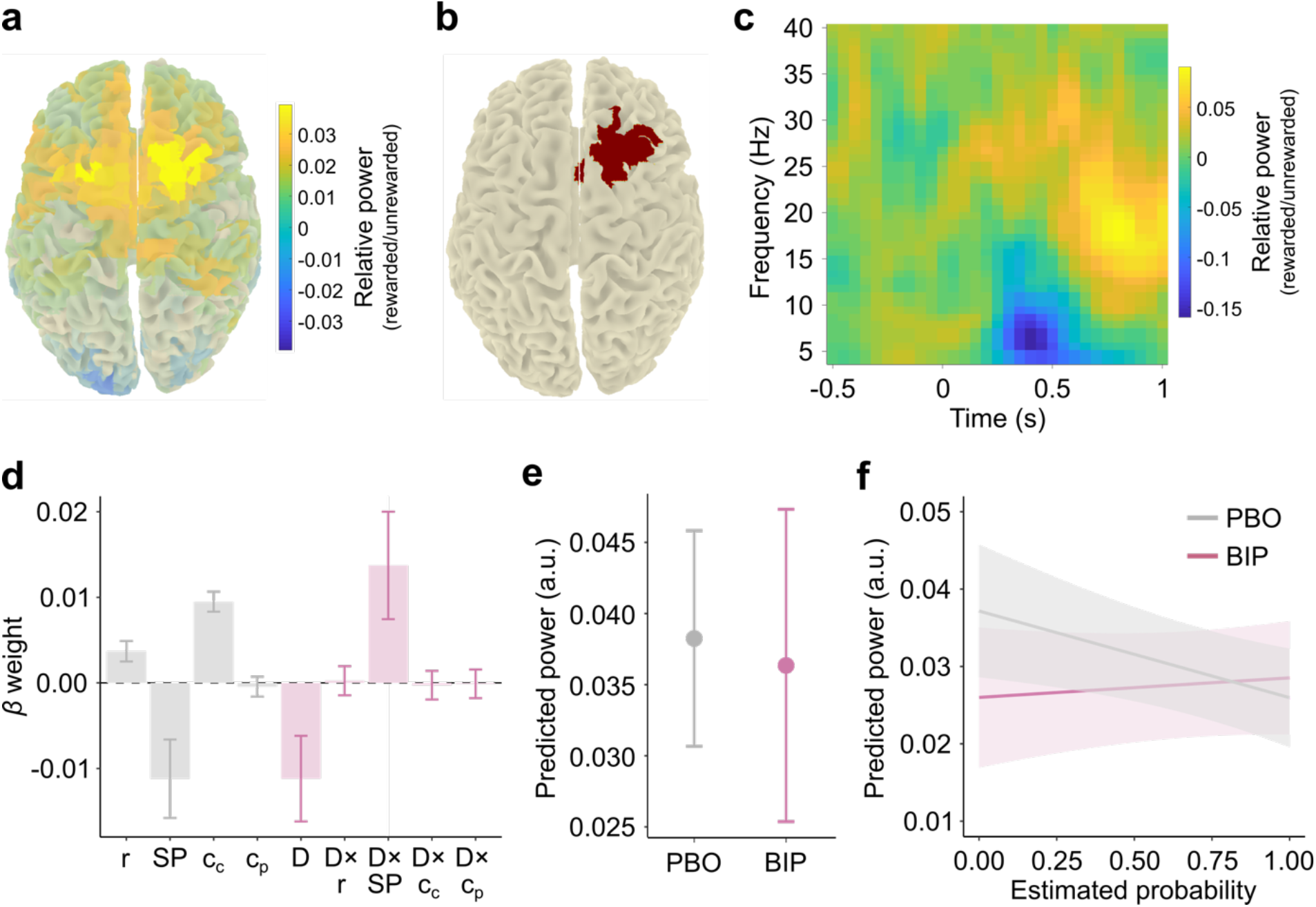
MEG analysis results for the volatile phase of the learning task. **a** Average high-beta-band power in rewarded relative to unrewarded trials (With higher power marked in yellow, lower power in blue). **b** ROI based on **a** (and supplementary Fig. S8d) in right dorsolateral prefrontal cortex. **c** Time-frequency representation (TFR) for rewarded relative to unrewarded trials, averaged within the ROI (colour conventions as in a) **d** Effects of task parameters (outcome r and subjective reward probability estimate SP), motor responses (current and previous choice side, c_c_ and c_p_), drug D, and interactions with drug on high-beta-band power within the ROI. Beyond significant task and motor effects, biperiden lead to an overall decrease in beta power and it modulated the effect of probability on betas power. **e** Main effect of drug on high-beta power. Beta power is significantly reduced under biperiden (BIP, pink) compared to placebo (PBO, grey). **f** Modelled beta power (based on the mixed-effects regressions) as a function of subjects’ estimated reward probability, separately for the biperiden and placebo session. Post-hoc tests indicate that the significant negative modulation of beta power by estimated reward probability under placebo (PBO, grey) is abolished under biperiden (BIP, pink). Solid lines represent mean, shaded areas 95 % CI.

In the ROI, we first observed the hallmarks of a prediction error signal (Rutledge et al., 2010): Under placebo, beta power increased for positive relative to negative outcomes (by definition of the rewarded vs. unrewarded ROI selection), and scaled negatively with estimated reward probability (*b* = -1.12×10^-2^, *SE* = 4.58×10^-3^, *t*(12270) = -2.45, *p* < .014, Fig. 4d; see supplementary Table S12 for full results). Biperiden modulated high-beta power in our lateral prefrontal ROI. First, we observed a main effect of drug: The overall high-beta response to feedback was reduced under biperiden compared to placebo (*b* = -1.12×10^-2^, *SE* = 5.01×10^-3^, *t*(603.3) = -2.23, *p* = .026, Fig. 4d, e). More importantly, biperiden also attenuated the prefrontal representation of the prediction error. In particular, biperiden impaired the estimated reward probability component of the prediction error, as evident from an interaction between drug and probability estimate (*b* = 1.37×10^-2^, *SE* = 6.28×10^-3^, *t*(11410) = 2.12, *p* = .029, Fig. 4f). To follow up on this interaction effect, we performed a post-hoc analysis for the biperiden condition only. Under biperiden there was no longer a significant main effect of estimated probability, indicating that the interaction effect reflects an elimination of the modulation of high-beta power by estimated probability under biperiden (see supplementary Table S13 for results). This reveals that, upon reward delivery, biperiden impaired a crucial computational step necessary for appropriately updating value estimates for future choice. The drug effect seems to be specific for the prediction component of the prediction error (i.e. probability tracking), since there was no significant interaction between drug and outcome (*p* = .880). In addition to our main task parameters of interest, we also included the choice (left vs. right) on the current and previous trial in our model to account for the well-known effects of movement-related variables on cortical beta power. Motor choice on the current (but not previous) trial was represented in prefrontal beta power (*b* = 9.49×10^-3^, *SE* = 1.17×10^-3^, *t*(13770) = 8.09, *p* < .001, Fig. 4d). This effect of choice was not affected by biperiden (*p* = .873), indicating that our drug effects on prediction error components are independent of effects on representations of motor actions. For comparison, we conducted the same analysis in the left dorsolateral prefrontal cortex. Here, motor choice both on the current and previous trial had a main effect on beta power (current choice side: *β* = -5.57×10^-3^, *SE* = 1.13×10^-3^, *t*(13760) = -4.92, *p* < .001; previous choice side: *β* = 2.72×10^-3^, *SE* = 1.13×10^-3^, *t*(13760) = 2.40, *p* = .016). However, we observed no effect of estimated probability in this ROI (see supplementary Table S14 for full results), nor a main effect of drug on beta power, indicating that both the signature of prediction error and the biperiden-induced (main and interaction) effects were specific for the right lateral prefrontal cortex.

Taken together, in keeping with previous single-cell recording studies in primates (Barraclough et al., 2004; Seo et al., 2007; Seo & Lee, 2009), these results indicate that lateral prefrontal high-beta power carries representations of outcomes, motor actions on the current (and previous) trial, and probability estimates inferred via reinforcement learning. Importantly, biperiden disrupted this prefrontal representation of estimated probability, indicating impaired computation of prediction errors.

## Discussion

Decision making requires flexibility, especially, when options or outcomes are uncertain. One approach to adapt behaviour involves increasing the learning rate in volatile environments (Behrens et al., 2007). Cholinergic transmission is crucial for learning and uncertainty processing (Everitt & Robbins, 1997; Hasselmo & Sarter, 2011) and has been suggested to govern the learning rate in reinforcement learning (Doya, 2002). Therefore, in the present study, we investigated the effects of biperiden, a muscarinic acetylcholine receptor antagonist, on decision making in healthy (male) participants in three scenarios: (i) under risk, when all attributes were explicitly presented, (ii) under uncertainty, when one attribute needed to be learnt, and, on top of learning, (iii) under volatility, characterized by frequent changes of stimulus-outcome contingencies. Biperiden affected decision making when option attributes needed to be learnt from experience, but had no effect on choice behaviour when all reward attributes were explicitly presented. More specifically, biperiden reduced sensitivity to the learnt reward probabilities selectively in the volatile phase of the learning task. Computational modelling revealed that this reduced sensitivity to Bayes-optimal reward probabilities was caused by impairments in the accurate estimation of implicit reward probabilities rather than by reduced reliance on learnt reward probability. Biperiden increased the learning rate for rewarded choices in the volatile phase in a maladaptive manner, such that participants’ probability estimates exhibited stronger deviations from the statistically optimal probability estimates compared to placebo. At the time of outcome presentation, high-beta power in lateral prefrontal cortex carried representations of both outcome and estimated outcome probability, the key terms of a prediction error signal. Importantly, the neural representation of estimated probability was abolished under biperiden. This presents a plausible neural mechanism of the maladaptive setting of learning rates observed at the behavioural level.

### Role of acetylcholine in volatile environments

Acetylcholine is widely distributed in the brain and known to influence behaviour such as attention, memory, and learning of cue-outcome associations (reviewed e.g. in Ananth et al., 2023). Theoretical proposals have suggested that acetylcholine may control the learning rate (Doya, 2002). This is indeed what we observed. Blockade of muscarinic receptors lead to a maladaptive increase in learning rate in the volatile phase of the learning task. This means that, under biperiden, participants put more weight on recent relative to more distant experience, leading to noisier, suboptimal estimation of reward probability. These data suggest a reduced integration over time, or even enhanced distractibility by current sensory (reward) information. This is in line with the observation that cholinergic antagonism with biperiden abolished post-error adjustments in both behaviour and sensory cortical areas (Danielmeier et al., 2015) and with findings of impaired serial reversal learning in a set-shifting task following depletion of cortical acetylcholine by lesions to the nucleus basalis magnocellularis (Cools & Arnsten, 2022; Robbins & Roberts, 2007). Similarly, Marshall et al. (2016) observed impaired adaptations to environmental changes in a probabilistic serial response time task under biperiden. In line with this, we found that the increase in learning rate was specific for the volatile phase which is also characterized by a higher degree of overall outcome surprise. This supports the hypothesis that participants tended to interpret probabilistic outcomes more readily as indicators for context changes, leading to noisier estimations of volatility (Marshall et al., 2016; Piray & Daw, 2021; Yu & Dayan, 2005).

Acetylcholine acting at M1 receptors enhances NMDA and GABA receptor function (Bessie Aramakis et al., 1997; Obermayer et al., 2017; Zwart et al., 2018). The balance between recurrent NMDA-mediated excitation and GABAergic feedback inhibition is a fundamental determinant in cortical circuit models of decision making (Wang, 2002; Wong & Wang, 2006). In line with this, human participants with higher concentrations of GABA relative to glutamate in the ventromedial prefrontal cortex were found to have a higher decision accuracy in a reward-guided choice task (Jocham et al., 2012; Kaiser et al., 2021). On these grounds, we had hypothesized that muscarinic antagonism would lead to impaired decision making in all tasks, beyond effects on learning and those induced by volatility manipulations. The apparent lack of a biperiden effect on choice behaviour in the gambling task and the stable phase of the learning task is therefore unexpected. We can only speculate about the reasons for this lack of effect. One possible reason is that our tasks were more difficult than those in earlier studies addressing related questions. In support of this, our participants used less multiplicative attribute integration (Farashahi et al., 2019; Scholl et al., 2014), and presented remarkably high learning rates in the learning task, indicative of suboptimal behaviour. Our drug effects on learning rates were specific to the volatile phase of the learning task. At first glance, this volatility-specific effect appears surprising given that acetylcholine is proposed to play a critical role in environments with known unreliability, often referred to as expected uncertainty. By contrast, uncertainty arising from unpredictable switches of context, such as reversal of cue-outcome-contingencies, is often labelled as unexpected uncertainty and has been ascribed to the neuromodulator noradrenaline (Avery et al., 2012; Yu & Dayan, 2005). By this definition, the gambling task and the stable phase of the learning task should be associated with expected uncertainty, whereas the volatile phase of the learning task involves elements of both expected and unexpected uncertainty. However, one notable prediction of this framework is that, under reduced cholinergic transmission, the degree of randomness in the environment is underestimated, which in turn should amplify the effect of unexpected outcomes - as these are then more likely to be taken as an indication that stimulus-outcome-contingencies have switched. Indeed, the authors refer to the acetylcholine-depleted state of their model as “hyper-distractible” (Yu & Dayan, 2005). In line with both this theoretical framework and our experimental results, Marshall et al. (2016) also observed more rapid updating of higher-order volatility estimates under biperiden. In this context, it is also worth noting that the mean squared prediction error was higher in the volatile compared to the stable environment. This might explain why biperiden specifically affected the volatile phase of the learning task, where the increased level of surprise in the environment under reduced cholinergic transmission is more readily interpreted as a change in outcome contingencies. While the only robust effect of biperiden was an increase in the learning rate for rewarded choices, we also observed numerical increases in the learning rate for unrewarded choices, greater choice stochasticity, and reduced weighting of probabilities relative to magnitudes. These effects were not credibly detected in our relatively large sample, suggesting that any influence of biperiden on these parameters is likely to be subtle. Nevertheless, small changes across multiple parameters might jointly contribute to the overall behavioural pattern observed under biperiden.

The observed effects of cholinergic antagonism are complemented by pharmacological studies increasing catecholaminergic transmission that found similar effects. In particular, the catecholamine reuptake inhibitor methylphenidate has been reported to increase learning rates in a volatile environment of a learning task similar to ours (Cook et al., 2019), suggestive of opposing actions of muscarinic acetylcholine and catecholaminergic transmission. For the dopaminergic system, there is evidence for a reciprocal antagonistic interaction between cholinergic M1 and dopaminergic D2 receptors at the behavioural (Brocks, 1999; Stanhope et al., 2001) and cellular level in the striatum (Di Chiara et al., 1994). In line with this, we recently observed that biperiden had effects on effort-based decision making that were opposite to those of the D2 receptor antagonist haloperidol (Erfanian Abdoust et al., 2024).

### Cholinergic modulation of prefrontal learning computations

We found that high-beta power in the right lateral prefrontal cortex was modulated by several task parameters of interest, including motor action, outcome, and estimated outcome probability. In particular, following outcome presentation, beta power was modulated in opposite directions by outcome and estimated probability (positive for outcome, negative for estimated probability). This is the key signature of a prediction error response. Importantly, biperiden abolished the prefrontal representation of estimated probability in high-beta power, thereby interfering with a key computation required for updating probability estimates for choices on future trials. This effect of biperiden therefore presents a candidate neural mechanism for the maladaptively changed learning rates. It is important to note that these value-related modulations are unlikely to be related to motor effects. MEG and EEG studies often report reductions in beta power around the time of movement onset, followed by later overshoot, the post-movement beta-rebound (Fischer et al., 2018; Pape & Siegel, 2016; Pfurtscheller et al., 1997; Rogge et al., 2022). However, firstly, these negative modulations are observed in regions substantially posterior to our region of interest. Secondly, even when sensorimotor beta power is modulated by motor action, orthogonal co-modulations by expected value have still been observed (Hunt et al., 2013). Thirdly, we used a regression-based approach to account for different sources of variance that jointly contribute to beta power. We first selected our region of interest based on a reward (rather than motor) response and included both learning-related parameters (outcome and estimated probability) and motor parameters (motor action on the current and previous trial) to our model. Accordingly, we observed orthogonal representations of choice, outcome, and estimated probability in high-beta power. These findings are in keeping with previous single-cell recording studies in primates that also evidenced multiplexed coding of choices and outcomes in lateral prefrontal cortex (Barraclough et al., 2004; Seo et al., 2007).

Previous literature has already indicated that beta power may reflect a number of cognitive parameters, in addition to motor-related variables. Post-movement decreases in beta power have been related to prediction errors and to updating of an internal model (Tan et al., 2014, 2016). Furthermore, beta power has also been demonstrated to reflect uncertainty measures (Palmer et al., 2019; Tan et al., 2016; Tzagarakis et al., 2015; Winkler et al., 2025) and to be reduced under conditions of higher predictability in a perceptual task (van Pelt et al., 2016). In addition to this, beta-band activity has also been linked to statistical learning (Bogaerts et al., 2020), increased proactive attentional control (van Driel et al., 2019), top-down signalling of (learned) predictions (Fries, 2015; Lee et al., 2013; Richter et al., 2018), and recruitment of task-relevant neural ensembles during decision making (Haegens et al., 2017). Notably, several previous studies reported that cortical beta oscillations are modulated by acetylcholine. Stimulation of the nucleus basalis magnocellularis in rats led to cortical desynchronization and increased high-frequency oscillations (Metherate et al., 1992). This effect appears to be largely mediated by muscarinic receptors, since administration of muscarinic antagonists in both humans and animal models, in the absence of a behavioural task, usually leads to a synchronized cortical state with increases in low-frequency oscillations and decreases of power in higher frequencies (Bakker et al., 2021; Dimpfel, 2005). It is noteworthy however that the exact pattern of results varies between studies (Kikuchi et al., 1999; Knott et al., 1997; Osipova et al., 2003) and may depend on the particular agent used. Scopolamine and atropine bind to all five muscarinic receptor subtypes with comparable affinity. In contrast, biperiden, while also binding to all five subtypes, has about tenfold higher affinity for the M1 receptor compared to the other subtypes (Bolden et al., 1992). Indeed, Dimpfel and colleagues (2005) highlighted that biperiden had effects on rat EEG that, while mostly consistent with those of scopolamine, in parts were in the opposite direction. Our finding that biperiden, over and above task-related effects, lead to a decrease in high-beta power is consistent with effects described by Bakker and colleagues (2021), who used the same dose of biperiden (4 mg) as in the present work. In addition to task-independent modulations of cortical oscillations, it has also been reported that increasing cholinergic tone by administration of physostigmine amplified the attentional modulation of alpha and beta oscillations in visual areas (Bauer et al., 2012). In general, the above findings dovetail with our findings of both a main drug effect (overall reduction of beta power) and a disruption of the modulation of beta power by estimated probability in lateral prefrontal cortex.

### Behavioural adaptions across tasks

While numerous studies have reported an increased learning rate during volatile relative to stable phases in humans (Behrens et al., 2007; Blain & Rutledge, 2020; Browning et al., 2015), we were surprised not to observe this pattern. Our data suggest that learning rate adjustments in response to volatility are less robust/consistent when the difficulty of learning is high, such as when the value difference between options or the average reward rate in the environment is low. In studies that observed learning rate adjustments, volatility has typically been more discernible, either in terms of reward probabilities (Behrens et al., 2007; Browning et al., 2015) or in terms of explicit cueing of phases (Blain & Rutledge, 2020; Massi et al., 2018). Both makes probability estimates less uncertain, and, more importantly, contingency switches easier to detect. This speculation also aligns with the interpretation of Cook et al. (2019), who hypothesized that two sources of information for learning might have interfered with learning rate adjustments in their study.

In addition to adjusting the learning rate, there is also evidence that the valuation strategy, i.e., the way in which values are computed, may vary with different levels of uncertainty (Farashahi et al., 2019). Although most commonly used models, such as prospect theory, assume that reward probability and magnitude are combined multiplicatively to estimate the options’ values (Bernoulli, 1954; Kahneman & Tversky, 1979), it has been suggested that agents also employ an additive strategy to combine information (Stewart, 2011), in particular under high uncertainty, e.g. when reward probabilities need to be learnt (Farashahi et al., 2019). We observed that, in the gambling task, participants used a mixture of both multiplicative and additive value construction, the latter with a strong weighting of reward probability. In contrast, in the learning task, there was no evidence for multiplicative integration and participants’ choices relied less on (estimated) reward probabilities relative to magnitudes compared to the gambling task. Thus, participants appear to adjust their decision strategy to be more flexible under conditions of higher uncertainty.

Taken together, our results suggest that muscarinic receptors play a key role in appropriately adapting learning rates and for prefrontal value computations in highly uncertain environments, when the demand for carefully calibrated adjustments is greatest.

## Materials and methods

### Ethics statement

All procedures were approved by the Ethics Committee of the Medical Faculty of the Heinrich Heine University Düsseldorf (reference 2018-211_1). The study was performed in compliance with the Code of Ethics of the World Medical Association (Declaration of Helsinki, 1975).

### Participants

Participants were recruited from the local student community of the Heinrich Heine University Düsseldorf, Germany. Participants signed a written informed consent prior to participation and received monetary compensation for their participation. Each experiment was run with healthy males who reported normal or corrected-to-normal vision. Due to the pharmacological challenge, recruited participants were extensively screened for medical exclusion criteria (see supplementary list S1). Additionally, participants were only included if they succeeded at both tasks in the screening session. For that reason, we set a performance criterion of expected value choices > 60 % for the gambling task and > 55 % for the learning task. In total, 43 participants aged between 18 and 35 years (mean age = 23.7 ± 3.1 years) took part in the study. Due to technical issues, MEG data of three participants had to be excluded leading to an inclusion of 40 participants in MEG analyses.

### General design

We pharmacologically manipulated the levels of acetylcholine in a double-blind, randomized, placebo-controlled, within-subjects design. Each participant completed four sessions: one screening session, two experimental sessions, which took place in the MEG, consisting of drug or placebo intake, and one MRI session. The screening session took place on a separate day prior to the pharmacological experimental sessions. After providing informed consent, we tested whether medical inclusion criteria were fulfilled and measured heart rate, blood pressure and the Beck’s Depression Inventory score (Beck et al., 1996). In addition, participants conducted the State-Trait Anxiety Inventory (STAI; Spielberger, 1983) and a modified version of the Edinburgh Handedness Inventory (Oldfield, 1971). Finally, participants performed the two behavioural tasks to assure that they exceeded pre-defined performance thresholds (choice of high expected value option > 60 % and > 55 % for the gambling and learning task, respectively). The two experimental sessions took place at University Hospital Düsseldorf. After over-night fasting and a standardized breakfast, participants received a single oral dose of biperiden on one day and a placebo on the other day 45 minutes prior to the MEG recording (Fig. 1a). Blood pressure, heart rate and mood (Bond and Lader Visual Analogue Scales; Bond & Lader, 1974) were measured after breakfast, before entering the MEG chamber, and at the end of the session. Before entering the MEG chamber, participants additionally conducted a modified trail-making test (Rodewald et al., 2012; part A). During MEG measurement, participants were seated on a chair inside a dimly lit, magnetically shielded MEG chamber. Each MEG measurement started with a 5 minutes eyes-open resting state task followed by the two behavioural paradigms with 500 trials of the gambling task (∼30 min) and 400 trials of the learning task (∼20 min). In a last and fourth session we recorded an anatomical MRI scan.

### MEG recording

We recorded neuromagnetic brain activity using a whole-head MEG system with 102 magnetometers and 204 planar gradiometers (Neuromag Elekta Oy). The sampling rate of the recordings was 1 kHz and MEG signals were filtered with a high-pass filter at 0.1 Hz and a low-pass filter at 330 Hz. To detect eye blinks and heartbeat, vertical and horizontal electrooculograms (EOG) and electrocardiograms (ECG) were recorded, respectively. Head position in the MEG helmet was registered at the beginning of each task using four head position indicator (HPI) coils attached to the scalp. HPI coil locations, head surface points from across the scalp and three anatomical fiducial locations (nasion, left and right pre-auricular points) were digitized using a Polhemus Isotrak before data acquisition. We acquired full-brain standard T1-weighted structural magnetic resonance images of each participant on a 3 T MRI scanner (Siemens MAGNETOM Trio A TIM System) using an MPRAGE (magnetization-prepared rapid gradient echo) imaging sequence (repetition time = 2000 ms, echo time = 2.45 ms) for source-localization of neuromagnetic activity.

### Pharmacological intervention

We administered the muscarinic acetylcholine receptor antagonist biperiden (4 mg). Typically, peak plasma concentrations are reached between 1 and 1.5 hours after oral administration of biperiden and the elimination half-time is about 18 to 24 hours (Brocks, 1999; Grimaldi et al., 1986). In view of this pharmacokinetic profile, MEG measurements started 45 minutes after drug administration and lasted approximately 1.5 hours. To ensure complete washout of the drug before the next session, the two experimental sessions were scheduled at least 6 days apart. After the final session, we tested participants’ awareness of the drug manipulation and found 35 participants to guess correctly post-hoc on which testing day they had the treatment (i.e. biperiden) vs. placebo pill with a certainty of 77.5 ± 3.9 (mean ± SEM) on a scale of 1 to 100.

### Behavioural tasks

The experiment was designed and presented using the PsychoPy software package (version 3.1.5; Peirce, 2007). Inside the MEG chamber, stimuli were presented on a projector (Panasonic PT-D7700E, screen dimensions: 43 cm × 31 cm) with a resolution of 1280 x 1024 pixels and a refresh rate of 60 Hz (viewing distance: 80 cm). For the behavioural tasks, participants responded bi-manually with their left and right index fingers on a custom-made button box with only two buttons available.

#### Gambling task

We implemented a gambling task in which participants had to decide between two options on each of 500 trials in order to maximize their reward. Each trial started with the presentation of a fixation dot (radius = 0.4 dva), for a pseudo-randomly selected duration of 1000 ms to 1400 ms to keep the participants’ attention to the centre of the screen. Afterwards, the two options (width = 0.6 dva, height = 1.36 dva) appeared on the left and right side (2.2 dva distance) of the screen until a response was made. The options were visualized as vertical bars, with the fill level indicating the reward magnitude and a numeric percentage below each bar indicating the reward probability (Fig. 1). The cumulative reward already earned was visualized by a progress bar at the bottom of the screen. Participants had 3000 ms to make a choice by pressing the left or right button, followed by the presentation of a frame around the chosen option for 400 ms to 600 ms. Next, the outcome of both options was presented for 500 ms by changing the colour of the bars to either green or red indicating whether the option was rewarded or not, respectively. The outcomes of both options were independent of each other. When no response was given in time, a warning was presented for 500 ms, urging participants to respond within the time frame. The task consisted of ten blocks of 50 trials. Participants were encouraged to rest between blocks as long as they needed, but had to wait for at least 10 s before the next block could be started.

#### Learning task

In the learning task, participants again had to choose between two options to maximize their reward. However, in contrast to the gambling task, not all relevant information of the options was explicitly presented, requiring participants to learn an implicit choice attribute. Here, the reward probability was implicit, similar to Farashahi et al. (2019). The trial structure of the gambling and learning tasks was similar: After the presentation of the fixation dot for 1000 ms to 1400 ms (radius = 0.4 dva), both options were presented as vertical bars on the left and right side of the screen until response (distance = 2.2 dva, width = 0.6 dva, height = 1.36 dva). The reward magnitude was explicitly presented as fill level of both bars; However, the reward probability was linked to the colour of the bars, requiring participants to learn which of the two colours was associated with a high reward probability. The high-probability colour had a reward probability of 0.7, while the other had a reward probability of 0.3. Participants had 3000 ms to make a choice by pressing the left or right button which was followed by a frame around the chosen option for 400 ms to 600 ms. Whether the chosen option was rewarded or not was indicated by a smiley or frowny, which was presented for 500 ms in the centre of the screen (radius = 0.67 dva). Note that the outcomes of the two options were dependent in the learning task, thus if one option was rewarded, the other was unrewarded. A rewarded choice led to an increase in the progress bar. When no response was given in time, a warning was presented for 500 ms. Volatility levels were manipulated during the task by changing reward contingencies over the course of the experiment, involving a stable and volatile phase, each lasting for 200 trials. While reward probabilities were fixed in the stable phase, probabilities reversed six times in the volatile phase: Successive reversals were separated by either 20 (three times) or 40 trials (four times). The order of stable and volatile phase was counterbalanced across participants. For each participant, the task structure remained the same for all sessions. The colour pairs indicating the reward probabilities were different for each session. Every 50 trials, participants had the possibility to rest as long as they needed.

### Behavioural analyses

#### Statistical analyses

We tested how the option attributes, and, for the learning task, the phase (stable vs. volatile) explained participants’ choice (left vs. right choices). Specifically, we investigated how biperiden changed the influence of the attributes on choices. We used logistic mixed-effects models using the lme4 package in R (R version 4.0.2 (2020-06-22); Bates et al., 2015). To account for within-subjects variability, we set the subjects’ ID as random effect for both tasks. The fixed effects, which account for the between-subjects variability, were dependent on the task: For the gambling task, we entered drug (biperiden vs. placebo), the reward magnitudes, and reward probabilities of both options and, their mean-centred product (i.e. expected value) and the previous choice side as between-subjects factor. Note that the expected value as calculated here recapitulates the multiplicative integration of reward magnitude and probability only. For the learning task, we additionally used the phase (stable vs. volatile) as between-subjects factor. However, since participants did not know the objective probabilities in the learning task, we used estimated probabilities from a Bayesian optimal learner, based on Behrens et al. (2007). The Bayesian optimal learner was set up in MATLAB (MATLAB Version R2016b, Massachusetts: The Mathworks Inc.) and run individually for each participant and session. In a model comparison approach, we tested whether the testing day improved the model fit. Additionally, we incorporated control measures (BL-VAS, TMT, heart rate and blood pressure) as fixed effects only when we observed significant drug effects on these measures. All *p*-values were based on asymptotic Wald tests.

#### Computational modelling

Computational modelling was performed in R (R version 4.0.2 (2020-06-22)).

#### Gambling task

To determine how participants used option attributes to shape a decision, we used a hybrid model, incorporating both additive and multiplicative value integration (Farashahi et al., 2019; Scholl et al., 2014). The subjective value *SV*_*i,t*_ for each option *i* at trial *t* was computed as follows:

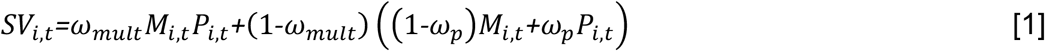

Where *M*_*i,t*_ is the reward magnitude, *P*_*i,t*_ is the reward probability, *ω*_*mult*_ is the degree of multiplicative relative to additive integration and *ω*_*P*_ is the degree of probability relative to magnitude weighting within the additive component. *ω*_*mult*_ would be either 1 or 0 if only the multiplicative or additive integration was used, respectively. Similarly, if participants only used the information of probability or magnitude, *ω*_*P*_ would be either 1 or 0, respectively. Magnitudes were scaled to values between 0.1 and 1.0 to allow for the comparison with reward probabilities. Based on the subjective values, the probability for choosing the left option *p*_*l,t*_ at trial *t* was generated using a softmax choice rule:

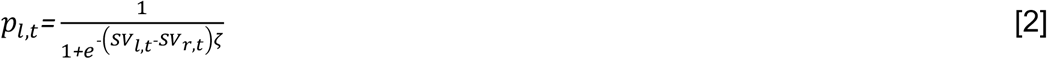

Where *SV*_*l,t*_ and *SV*_*r,t*_ are the subjective value of the left-side and right-side option, respectively, and *ζ* is the inverse temperature parameter capturing the stochasticity of action selection. We used hierarchical Bayesian estimation of group-level and subject-level parameters to incorporate the within-subjects design, similar to Swart et al. (2017). For individual-level parameters *x* group-level parameters *X* were used as priors *x* ∼ *N*(*X*, *σ*) and a half-Cauchy with a scale of 2 served as hyperprior for *σ* (Gelman, 2006). Weakly informative distributions were used as hyperpriors for *X*: *X*_*mult,P*_ ∼ *N*(0,2), *X* _*ζ*_ ∼ *N*(2,3). While *ω* _*mult*_ and *ω* _*P*_ were constrained between 0 and 1 using an inverse logit transform, *ζ* was positively bounded using an exponential transform. Initial parameter estimates were determined using an independent training dataset. The model allowed for a biperiden-induced shift *s*_*x*_ for all fitted parameters *x* with *x+δ*_*bip*_*s*_*x*_, where *δ*_*bip*_ is 0 or 1 for the placebo and biperiden session, respectively. The parameter shifts were unconstrained and *N*(0,3) served as hyperprior. In total, six free parameters were fitted. We performed Markov chain Monte Carlo (MCMC) sampling in RStan (RStan version 2.21.8, Stan Development Team, 2016). We used four Markov chains for sampling with 2500 iterations, including 500 warm-up iterations, per chain. Models successfully converged for a maximal potential scale reduction factor 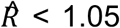 and after verifying convergence and diagnostic criteria, as provided by RStan. To verify if the model captures participants’ behaviour, we conducted posterior predictive checks by simulating 500 datasets based on the posterior distributions of subject-level parameters. We then compared and correlated simulated and real data (see supplementary Fig. S9-S11 and tables S15-S17).

#### Learning task

Computational modelling was similar for both tasks. Since our logistic mixed-effects regressions yielded that participants did not use the multiplicative strategy in the learning task, fitting of the hybrid model led to convergence issues. Thus, we fixed *ω* _*mult*_ at 0 for the computation of the subjective value, making the value integration additive only:

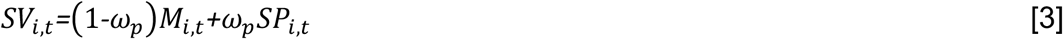

With the subjective probability *SP*_*i,t*_. The learning of reward probabilities was modelled using *Q*-learning. The probability estimate of the chosen colour *SP*_*c,t*_ was updated on each trial *t* via two separate learning rates for rewarded and unrewarded choices, *λ*_*r*_ and *λ* _*u*_, respectively:

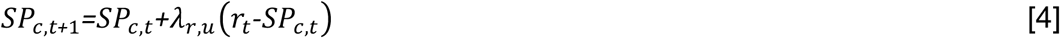

Where *r*_*t*_ reflects the outcome of the current trial and was either 1 or 0, depending on whether the choice was rewarded or not. The probability estimate of the unchosen colour *Sp*_*u,t*_ was dependent on the chosen colour:

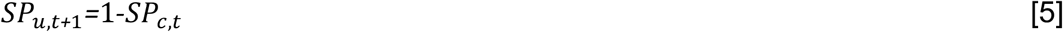

Again, parameters were estimated using Bayesian hierarchical modelling with a shift on each parameter in the biperiden session. Thus, eight free parameters were fitted. Learning rates were constrained between 0 and 1 via an inverse logit transform and *N*(0,2) was used as hyperprior. All other priors, hyperpriors and transformations were defined in the same way as in the gambling task.

### Data preprocessing

MEG data preprocessing was performed in Python (version 3.9.12), using MNE-Python (v.1.0.2; Gramfort et al., 2013) and custom written scripts. First, power line artifacts were removed using the ZapLine algorithm (de Cheveigné, 2020). Next, we marked bad channels based on visual inspection. All following preprocessing steps were performed on epoched data. We created epochs from -1000 ms to 1500 ms relative to feedback onset. In order to remove EOG- and ECG-related components, we conducted independent component analysis (ICA). We then excluded bad epochs based on visual inspection. Last, bad channels were interpolated.

### Source reconstruction

Source reconstruction was conducted in MATLAB using the FieldTrip toolbox (version 20191127; Oostenveld et al., 2011). We generated a single-shell head model for each participant based on their individual anatomical MRI scan. Sources were reconstructed for a cortical surface grid with 567 points aligned to Montreal Neurological Institute (MNI) space. For source reconstruction, we used a linear constrained minimum variance (LCMV) beamformer with the regularization parameter *λ* set to 5 %.

### MEG analyses

#### Time-domain analysis

Time-domain analysis was performed in FieldTrip. For the following analysis, we used frontocentral gradiometers with the following preprocessing settings: Data were demeaned and a band-pass filter of 1 – 40 Hz was applied using a windowed-sinc finite impulse response (FIRWS) filter. Additionally, padding of 3 s was used. A baseline correction using the interval from -0.1 to 0 s from feedback onset was applied.

### Time-frequency analysis

Time-frequency analysis was performed in FieldTrip. We used a Morlet wavelet transformation, with the wavelet width scaled linearly from 3 to 12 cycles. The analysis was performed over a time window from -0.5 to 1.5 s relative to feedback onset, sampled in 50 ms steps. Frequencies of interest ranged from 1 to 40 Hz. After the transformation, trials were truncated at 1 s, as the subsequent trial began at this time. Baseline correction was applied using the mean power of each trial.

#### Statistical analyses

We tested how the prediction error, decomposed into prediction and outcome components, and the motor action resulting from button presses explained high-beta band power. Specifically, we examined whether biperiden modulated the influence of prediction or outcome on power. To this end, we fitted mixed-effects regressions using the lme4 package in R including a treatment contrast. With this, the main effects of the independent variables reflect their influence on high-beta band power in the placebo condition, while the interaction between drug ID and the variables capture the biperiden-induced modulation of this effect. The dependent variable was the average high-beta band power within the 0 – 1 s time window after feedback onset. To account for within-subject variability, we included random intercepts and random slopes for drug ID by subject. Fixed effects, capturing between-subject variability, included subjective probability estimates derived from the Bayesian hierarchical model, current outcome (reward vs. no reward), current choice side and previous choice side. All *p*-values were based on asymptotic Wald tests.

## Code and data availability

All analysis code and behavioural datasets generated and analyzed during the current study are publicly available at: .

## Acknowledgments

The work was supported by a European Research Council grant (ERC-CoG 771432) to GJ. Computational infrastructure and support were provided by the Centre for Information and Media Technology at Heinrich Heine University Düsseldorf. We thank the MRI Core Facility of the University Hospital Düsseldorf, particularly Erika Rädisch and Eric Bechler, for help with the MRI recordings. We thank Hanin Alejel, Ana Antonia Dias Maile, Helena El Kholy, Judith Geusen, Paul Höchter, Marlene Hüsken, Christina Kalinicenko, Joshua Saal, Kouta Sasaki, Georg Schäfer, and Helena Schmidt for their support during data acquisition. The Bayesian optimal learner is a slightly modified version of MATLAB code kindly provided by Tim Behrens and James Whittington.

## Notes

### Competing Interest Statement

The authors have declared no competing interest.

### Summary of Updates

Manuscript and Supplemental files updated now including MEG data (analysis)

