## Supplementary Material for "Cholinergic modulation of reinforcement learning and prefrontal value computations under uncertainty"

### 1. Raw behaviour

Plotting the raw choice data as a function of the separate choice attributes illustrates that choice behaviour in both tasks was driven by the highest expected value (EV), which is the product of reward magnitude and probability, (figure S1, left side), by reward probabilities (figure S1, middle) and by reward magnitudes (figure S1, right side). Interestingly, in the gambling task, choices were only guided by magnitude information if the magnitude difference was large, for small magnitude differences participants opted for the other option (figure S1).

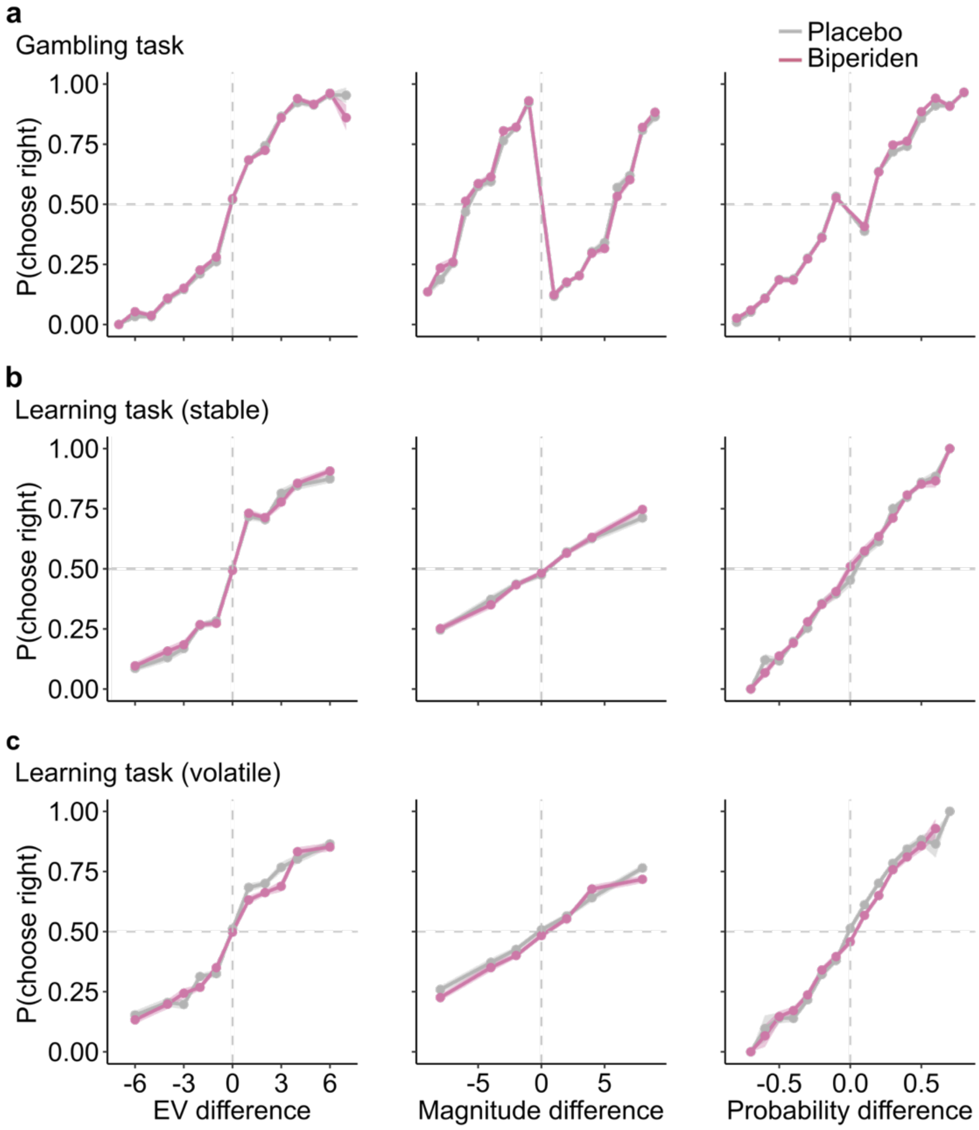

**Figure S1. Participants' choice behaviour.** Influence of task parameters on choice in the gambling task (**a**) and in both phases of the learning task (stable: **b**, volatile: **c**). Left: Probability of right-side choice as a function of difference in expected value (EV) between the two options. Note, that the shown EV is not mean-centred and, thus, is highly correlated with reward magnitude and probability. Middle: Probability of right-side choice as a function of difference in reward magnitude between the two options. Right: Probability of right-side choice as a function of difference in reward probability between the two options. For the learning task, reward probabilities were estimated using the Bayesian optimal learner. Choice behaviour in the biperiden session (pink) and the placebo session (grey) is shown. Solid lines represent mean, shaded areas SEM across participants.

### 2. Behaviour: Logistic mixed-effects regressions

Full results of the logistic mixed-effects regressions for the gambling task (Table S1) and the learning task (Table S2).

**Table S1: Results from logistic mixed-effects regressions of the gambling task.**

| | ***β*** | **SE** | ***z*** | ***p*** |
|---|---|---|---|---|
| Intercept | 0.03 | 0.03 | 0.86 | .392 |
| Drug | 0.03 | 0.02 | 1.37 | .172 |
| *Probability* | *2.90* | *0.03* | *95.78* | *< .001* |
| *Magnitude* | *1.58* | *0.02* | *73.66* | *< .001* |
| *EV* | *0.50* | *0.02* | *30.05* | *< .001* |
| *Repetition bias* | *-0.07* | *0.03* | *-2.64* | *.008* |
| Drug x probability | 0.00 | 0.03 | 0.12 | .902 |
| Drug x magnitude | -0.03 | 0.02 | -1.55 | .121 |
| Drug x EV | -0.02 | 0.02 | -1.02 | .310 |
| Drug x bias | -0.02 | 0.03 | -0.82 | .414 |

**Table S2: Results from logistic mixed-effects regressions of the learning task.**

| | ***β*** | **SE** | ***z*** | ***p*** |
|---|---|---|---|---|
| Intercept | -0.04 | 0.02 | -1.78 | .076 |
| Drug | -0.01 | 0.01 | -0.78 | .434 |
| *Probability* | *1.27* | *0.02* | *80.28* | *< .001* |
| *Magnitude* | *0.75* | *0.01* | *51.45* | *< .001* |
| *EV* | *-0.04* | *0.02* | *-2.31* | *.021* |
| *Repetition bias* | *0.10* | *0.01* | *7.86* | *< .001* |
| Phase | -0.01 | 0.01 | -0.63 | .529 |
| *Drug x probability* | *-0.05* | *0.02* | *-3.09* | *.002* |
| Drug x magnitude | 0.01 | 0.01 | 0.60 | .546 |
| Drug x EV | 0.02 | 0.02 | 1.39 | .164 |
| Drug x bias | 0.01 | 0.01 | 0.49 | .625 |
| *Phase x probability* | *-0.08* | *0.02* | *-4.97* | *< .001* |
| Phase x magnitude | 0.01 | 0.01 | 0.92 | .359 |
| Phase x EV | 0.00 | 0.02 | 0.31 | .757 |
| Phase x bias | -0.01 | 0.01 | -0.67 | .502 |
| *Drug x phase x probability* | *0.04* | *0.02* | *2.29* | *.022* |
| Drug x phase x magnitude | 0.01 | 0.01 | 0.38 | .706 |
| Drug x phase x EV | 0.01 | 0.02 | 0.52 | .604 |
| Drug x phase x bias | 0.02 | 0.01 | 1.60 | .110 |

### 3. Control measurements

During each session, we acquired the participants' mood, using the Bond and Lader Visual Analogue Scales (BL VAS), and blood pressure as control measurements at three time points: before drug intake (T1), before the MEG measurement (T2) and after the MEG measurement (T3). Before the MEG measurement, we additionally measured participants' executive functions using a trail making test. To control for biperiden effects on these measures, we applied linear mixed-effects regressions.

#### 3.1. Bond and Lader Visual Analogue Scales

Under biperiden, alertness, calmness, and contentedness were significantly decreased at T3 (see Table S3-S5, Figure S2).

**Table S3: Linear mixed-effects results for the alertness score.**

| | ***β*** | **SE** | ***df*** | ***t*** | ***p*** |
|---|---|---|---|---|---|
| *Intercept* | *7.90* | *0.23* | *63.31* | *34.22* | *< .001* |
| Drug | 0.13 | 0.16 | 209.01 | 0.85 | .396 |
| T2 | -0.02 | 0.16 | 209.01 | -0.10 | .917 |
| T3 | -0.08 | 0.16 | 209.01 | -0.54 | .592 |
| Drug x T2 | -0.06 | 0.22 | 209.01 | -0.29 | .776 |
| *Drug x T3* | *-1.23* | *0.22* | *209.03* | *-5.55* | *< .001* |

**Table S4: Linear mixed-effects results for the calmness score.**

| | ***β*** | **SE** | ***df*** | ***t*** | ***p*** |
|---|---|---|---|---|---|
| *Intercept* | *7.75* | *0.28* | *97.87* | *27.77* | *< .001* |
| Drug | -0.06 | 0.26 | 208.82 | -0.24 | .815 |
| T2 | -0.16 | 0.26 | 208.82 | -0.63 | .531 |
| T3 | -0.18 | 0.26 | 208.82 | -0.71 | .479 |
| Drug x T2 | 0.07 | 0.37 | 208.75 | 0.19 | .848 |
| *Drug x T3* | *-0.84* | *0.37* | *208.75* | *-2.29* | *.023* |

**Table S5: Linear mixed-effects results for the contentedness score.**

| | ***β*** | **SE** | ***df*** | ***t*** | ***p*** |
|---|---|---|---|---|---|
| *Intercept* | *8.53* | *0.20* | *67.45* | *43.31* | *< .001* |
| Drug | -0.03 | 0.14 | 209.07 | -0.18 | .858 |
| T2 | -0.16 | 0.14 | 209.07 | -1.17 | .244 |
| T3 | -0.14 | 0.14 | 209.07 | -1.00 | .316 |
| Drug x T2 | -0.02 | 0.20 | 209.04 | -0.11 | .914 |
| *Drug x T3* | *-0.55* | *0.20* | *209.04* | *-2.79* | *.006* |

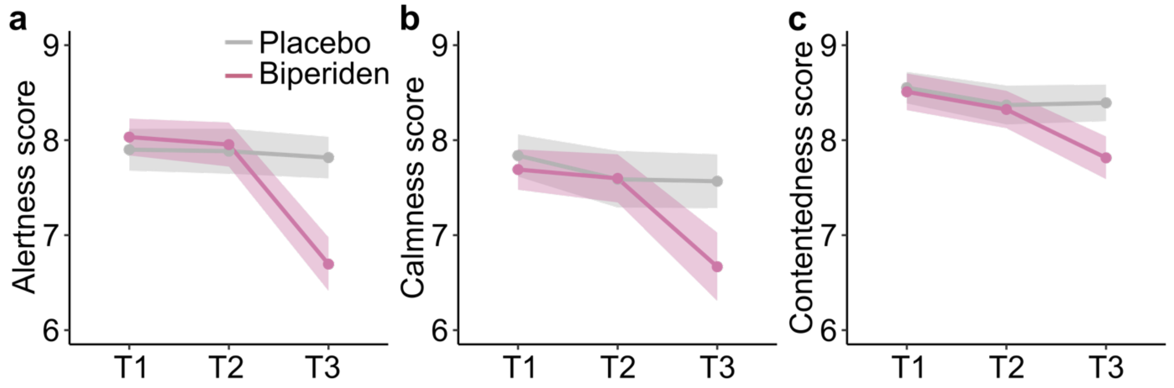

**Figure S2: Results of the BL VAS.** Scores for **a** alertness, **b** calmness, and **c** contentedness were acquired before drug intake (T1), before MEG measurement (T2), and after MEG measurement (T3) in both the placebo session (grey) and biperiden session (pink). All three measures were reduced at T3 under biperiden. Solid lines represent mean, shaded areas SEM across participants.

#### 3.2. Blood pressure

There were no significant effects of biperiden on systolic and diastolic blood pressure. However, the heart rate was decreased at T3 under biperiden (see Table S6-S8, Figure S3).

**Table S6: Linear mixed-effects results for the diastolic blood pressure.**

| | *β* | SE | *df* | *t* | *p* |
|---|---|---|---|---|---|
| *Intercept* | *124.48* | *1.69* | *92.78* | *73.76* | *< .001* |
| Drug | -2.69 | 1.53 | 208.04 | -1.76 | .080 |
| *T2* | *-3.64* | *1.52* | *208.21* | *-2.39* | *.018* |
| T3 | -0.62 | 1.52 | 208.21 | -0.41 | .684 |
| Drug x T2 | 3.41 | 2.15 | 208.04 | 1.59 | .114 |
| Drug x T3 | -3.57 | 2.15 | 208.04 | -1.66 | .099 |

**Table S7: Linear mixed-effects results for the systolic blood pressure.**

| | *β* | SE | *df* | *t* | *p* |
|---|---|---|---|---|---|
| *Intercept* | *78.29* | *1.31* | *87.14* | *59.82* | *< .001* |
| Drug | -1.62 | 1.14 | 208.07 | -1.42 | .158 |
| *T2* | *-5.31* | *1.14* | *208.22* | *-4.67* | *< .001* |
| *T3* | *-2.47* | *1.14* | *208.22* | *-2.17* | *.031* |
| Drug x T2 | 1.27 | 1.61 | 208.07 | 0.79 | .430 |
| Drug x T3 | -2.45 | 1.61 | 208.07 | -1.53 | .129 |

**Table S8: Linear mixed-effects results for the heart rate.**

| | **β** | **SE** | ***df*** | ***t*** | ***p*** |
|---|---|---|---|---|---|
| *Intercept* | *72.49* | *1.60* | *69.72* | *45.25* | *< .001* |
| Drug | 1.83 | 1.19 | 208.05 | 1.55 | .123 |
| T2 | -2.13 | 1.18 | 208.15 | -1.81 | .072 |
| *T3* | *-11.83* | *1.18* | *208.15* | *-10.03* | *< .001* |
| Drug x T2 | -3.07 | 1.67 | 208.05 | -1.84 | .067 |
| *Drug x T3* | *-8.97* | *1.67* | *208.05* | *-5.39* | *< .001* |

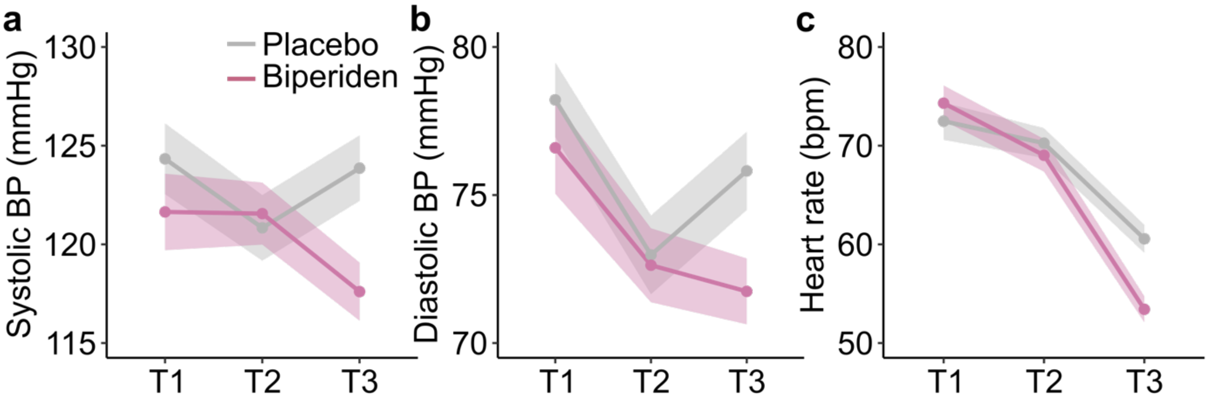

**Figure S3: Results of the measurement of the blood pressure.** Before drug intake (T1), before MEG measurement (T2), and after MEG measurement (T3) we measured **a** systolic blood pressure (BP), **b** diastolic BP, and **c** heart rate in both placebo session (grey) and biperiden session (pink). Biperiden significantly reduced heart rate at T3. Solid lines represent mean, shaded areas SEM across participants.

### 3.3. Trail making test

Participants' timing during the trail making test was not affected by biperiden ($\beta$ = -0.03, *SEM* = 0.84, $t_{214}$ = 0.037, $p$ = .971; figure S4).

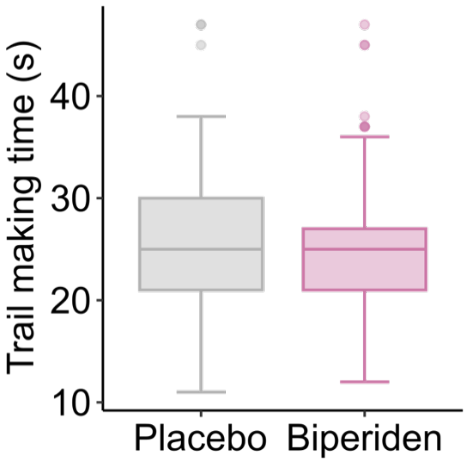

**Figure S4: Results of the trail making test before MEG measurement (T2).** Trail making time was acquired in the placebo session (grey) and in the biperiden session (pink).

### 3.4. Control for confounding effects in logistic mixed-effects regressions

After biperiden administration, we observed a significantly decreased sensitivity to estimated reward probabilities in the learning task, specifically in the volatile phase. To control if this biperiden effect is independent of other measures, we conducted logistic mixed-effects regressions for the volatile phase with several control measures (heart rate, alertness, calmness, and contentedness quantified by the BL VAS, order of stable and volatile phase, session days and order of medication).

**Table S9: Logistic mixed-effects results of the interaction effect of medication and estimated probability difference in the volatile phase considering control measures.**

| **Control measure** | ***β*** | **SE** | ***z*** | ***p*** |
|---|---|---|---|---|
| *Heart rate* | *-0.12* | *0.03* | *-4.42* | *< .001* |
| *Alertness* | *-0.06* | *0.03* | *-2.27* | *.023* |
| *Calmness* | *-0.08* | *0.03* | *-2.95* | *.003* |
| *Contentedness* | *-0.07* | *0.03* | *-2.76* | *.006* |
| *Phase order* | *-0.08* | *0.03* | *-3.33* | *< .001* |
| *Session days* | *-0.08* | *0.03* | *-3.35* | *< .001* |
| *Order of medication* | *-0.08* | *0.03* | *-3.32* | *< .001* |

### 4. Bayesian hierarchical models: Comparison between tasks

#### Results S1: Comparison of choice strategies within and across tasks

Behaviour under risk and uncertainty can be adapted in different ways. One approach is to adjust information integration between tasks (Farashahi et al., 2019), another approach is to increase the learning rate when contingencies change frequently (Behrens et al., 2007). To capture both possible approaches, we set up Bayesian hierarchical models. To foreshadow the findings, using these models, we found adaptations in attribute integration between tasks (gambling vs. learning task), but no adaptation within the learning task (stable vs. volatile).

In the models of both the gambling and the learning task, value can be constructed either in an additive or multiplicative fashion, or using a combination of both, which we call a hybrid strategy. Note that the purely additive and multiplicative models are nested within this hybrid model - they are special cases with values of $\omega_{mult}$ of either 0 or 1, respectively (see equations 1-5 for details). For the gambling task, we fitted a hybrid model, comprising both additive and multiplicative integration of option attributes, and softmax action selection. For the learning task, we used a similar model and added *Q*-learning with a delta update rule with learning rates $\lambda_r$ and $\lambda_u$, for rewarded and unrewarded choices, to capture how participants tracked probabilities. Because our logistic mixed-effects regressions (see Fig. 2 and Table S2) indicated that, in the learning task, multiplicative integration did not play a role in guiding participants' choices (resulting in convergence issues in the hybrid models), we allowed only additive value construction here (see methods for details). Stable and volatile phases were fitted separately, because the ultimate aim was to detail the phase-specific biperiden effects observed in the logistic mixed-effects regressions.

In line with the results from logistic mixed-effects regressions, in the gambling task, where both option attributes were explicitly presented, participants used both additive and multiplicative integration. Within the hybrid model, the parameter $\omega_{mult}$ was larger than 0 but below 1, which indicates a mixture of both strategies ($\omega_{mult}$: $HDI_{mdn}$ = 0.38, $HDI_{.95}$ = [0.25, 0.52], Fig. S5a). Reward probabilities were weighted more strongly than magnitudes ($\omega_P$: $HDI_{mdn}$ = 0.81, $HDI_{.95}$ = [0.75, 0.87], Fig. S5b). Similarly, in the learning task, participants also focused slightly more on (inferred) reward probabilities than on magnitude information (stable: $\omega_P$: $HDI_{mdn}$ = 0.59, $HDI_{.95}$ = [0.50, 0.66]; volatile: $\omega_P$: $HDI_{mdn}$ = 0.59, $HDI_{.95}$ = [0.51, 0.67], Fig. S5b), but relative probability weighting was much lower compared to the gambling task. There was no credible difference in relative attribute weighting between stable and volatile phase (Fig. S5b).

In the learning task, we observed that learning rates did not differ credibly between the stable and volatile phase, which was unexpected given earlier reports on learning rate adjustments (Behrens *et al.*, 2007; Browning *et al.*, 2015; Blain and Rutledge, 2020). In both phases, the learning rate for rewarded choices was higher than for unrewarded choices (stable: $\lambda_r$: $HDI_{mdn}$ = 0.60, $HDI_{.95}$ = [0.52, 0.68]; $\lambda_u$ = 0.25, $HDI_{.95}$ = [0.21, 0.30], Fig. 3C, volatile: $\lambda_r$: $HDI_{mdn}$ = 0.58, $HDI_{.95}$ = [0.52, 0.64]; $\lambda_u$: $HDI_{mdn}$ = 0.26, $HDI_{.95}$ = [0.22, 0.31], Fig. S5d).

Furthermore, as can be expected from the more uncertainty-laden value estimates in the learning task, choice stochasticity was increased compared to the gambling task. The softmax inverse temperature $\zeta$ was lower in the learning task compared to the gambling task, irrespective of stable and volatile phases (gambling task: $\zeta$: $HDI_{mdn}$ = 14.70, $HDI_{.95}$ = [12.92, 16.53]; learning task, stable: $HDI_{mdn}$ = 5.58, $HDI_{.95}$ = [4.80, 6.47], learning task, volatile: $\zeta$: $HDI_{mdn}$ = 5.60, $HDI_{.95}$ = [4.83, 6.40], Fig. S5e). Thus, participants were more deterministic in the gambling task than in the learning task, indicating that they were more sensitive to the estimated value of the options.

Although the fitted model parameters did not differ between the stable and volatile phase, we observed that the mean squared prediction error (MSPE) was significantly higher in the volatile compared to the stable phase ($\Delta(MSPE_{vol} - MSPE_{stab})$ = $2.48 \cdot 10^{-2} \pm 0.05 \cdot 10^{-2}$, mean ± SEM, $t(42) = 8.22$, $p < .001$, Cohen's $d = 1.25$, Fig. S5f). This reflects that, overall, participants experienced a higher degree of outcome-related surprise in the volatile phase.

In sum, when both option attributes were explicitly provided, participants used a hybrid attribute integration, consisting of mostly additive but also multiplicative integration. When learning was involved, participants showed no difference in decision strategies for stable versus volatile phases.

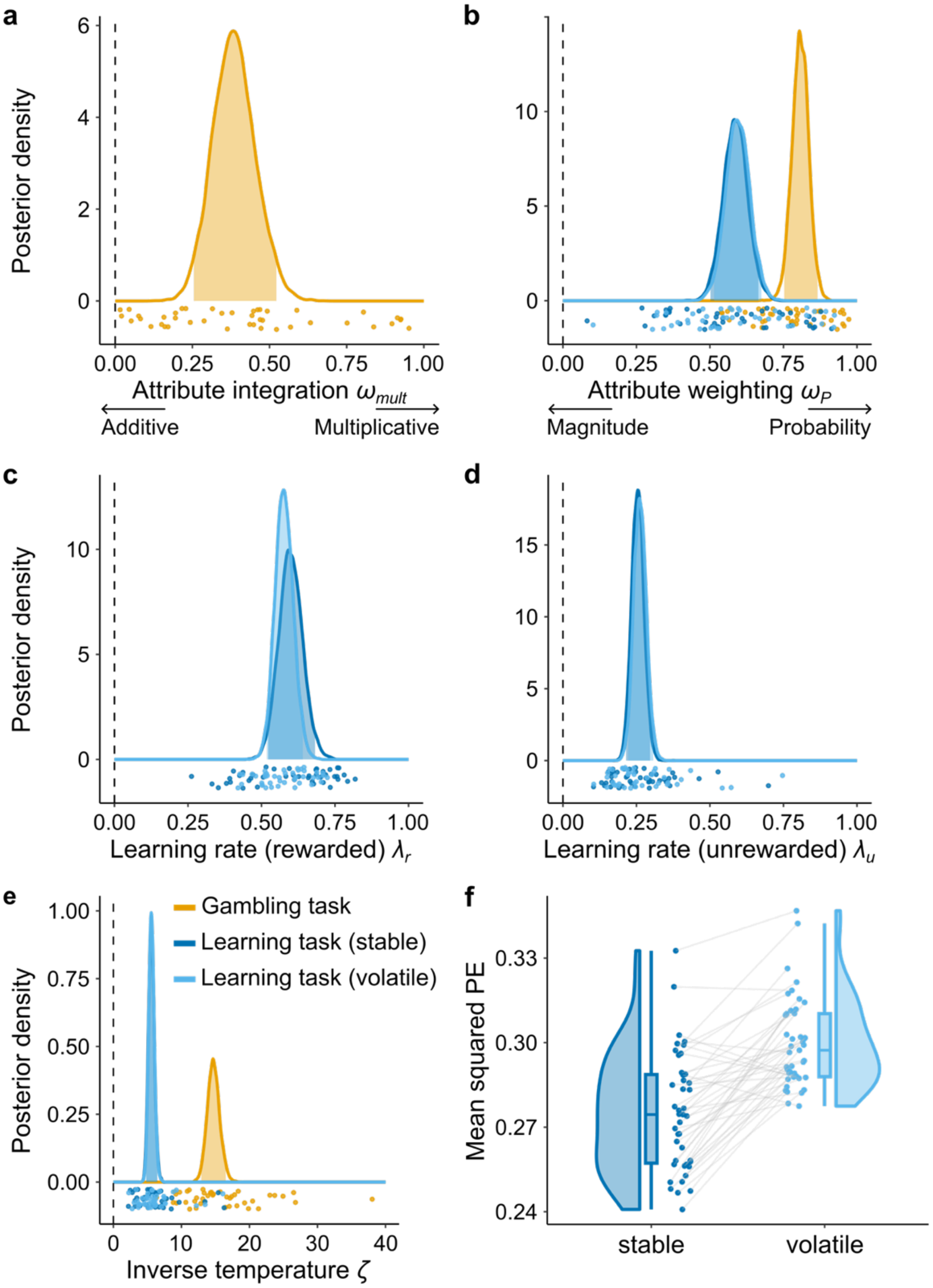

**Figure S5. Parameter fits of the Bayesian hierarchical model. a-e** Posterior densities for the gambling task (orange) and for the learning task (stable: dark blue, volatile: light blue), irrespective of drug effects. Shaded areas represent the 95 %-HDI of the posterior distribution, points are single-subject means. **a** Strategy for value construction $\omega_{mult}$. A purely multiplicative integration corresponds to $\omega_{mult}$ of 1, whereas $\omega_{mult}$ of 0 reflects a purely additive integration of option attributes. For the learning task, $\omega_{mult}$ was fixed at 0 (additive strategy). **b** Weighting of reward probability relative to magnitude $\omega_P$, where decisions based either entirely on probabilities or on magnitudes are reflected by values for $\omega_P$ of 1 or 0, respectively. In the gambling task, participants weighted reward probabilities more strongly than in the learning task. **c, d** Learning rates $\lambda_r$ for rewarded and $\lambda_u$ for unrewarded choices in the learning task. **e** Softmax inverse temperature $\zeta$. The higher the inverse temperature, the more deterministic the choice behaviour. Choice behaviour in the gambling task was more deterministic compared to the learning task. **f** Mean squared prediction error (PE) in the stable and volatile phase of the learning task. Points reflect individual participants' fit. In the volatile phase, the mean squared PE is significantly higher, indicating a higher degree of overall surprise.

### 5. Bayesian hierarchical models: Biperiden-induced shifts

#### 5.1. Gambling task

For the gambling task, we found no credible effect of biperiden-specific shifts in the Bayesian hierarchical models (Figure S6, Table S10).

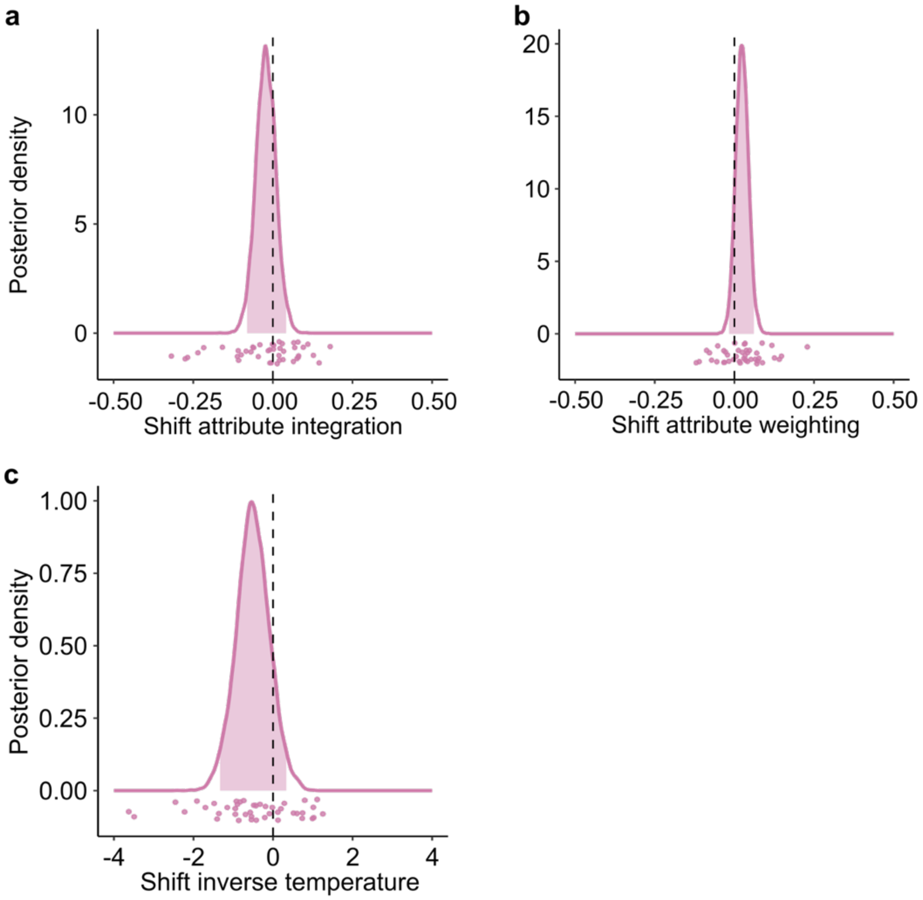

**Figure S6. Biperiden-induced shifts in the gambling task from the Bayesian hierarchical model.** Density of posterior predictive distributions of the biperiden-specific shift in **a** attribute integration $s_{\omega mult}$, **b** attribute weighting $s_{\omega P}$, and **c** inverse temperature $s_{\zeta}$. Positive shifts represent an increase under biperiden relative to placebo. Shaded areas represent the 95 %-HDI of the posterior predictive distribution and points single-subject means.

**Table S10. Group-level parameter estimates of the gambling task.** Median (Mdn), standard deviation (SD), and lower and upper bounds of the 95 %-HDI interval are given. The model consisted of estimates for the value construction $\omega_{mult}$, the attribute weighting $\omega_P$, the inverse temperature $\zeta$, and the corresponding biperiden-specific shifts on these parameters $s_{\omega mult}$, $s_{\omega P}$, and $s_{\zeta}$.

| **Parameter** | **Mdn** | **SD** | **2.5 %** | **97.5 %** |
|---|---|---|---|---|
| $\omega_{mult}$ | *0.38* | *0.07* | *0.25* | *0.52* |
| $\omega_P$ | *0.81* | *0.03* | *0.75* | *0.87* |
| $\zeta$ | *14.70* | *0.92* | *12.92* | *16.53* |
| $s_{\omega mult}$ | -0.02 | 0.03 | -0.08 | 0.04 |
| $s_{\omega P}$ | 0.02 | 0.02 | -0.02 | 0.06 |
| $s_{\zeta}$ | -0.51 | 0.42 | -1.33 | 0.33 |

5.2. Learning task: stable phase

For the stable phase of the learning task, we found no credible effect of biperiden-specific shifts in the Bayesian hierarchical models (Figure S7, Table S11).

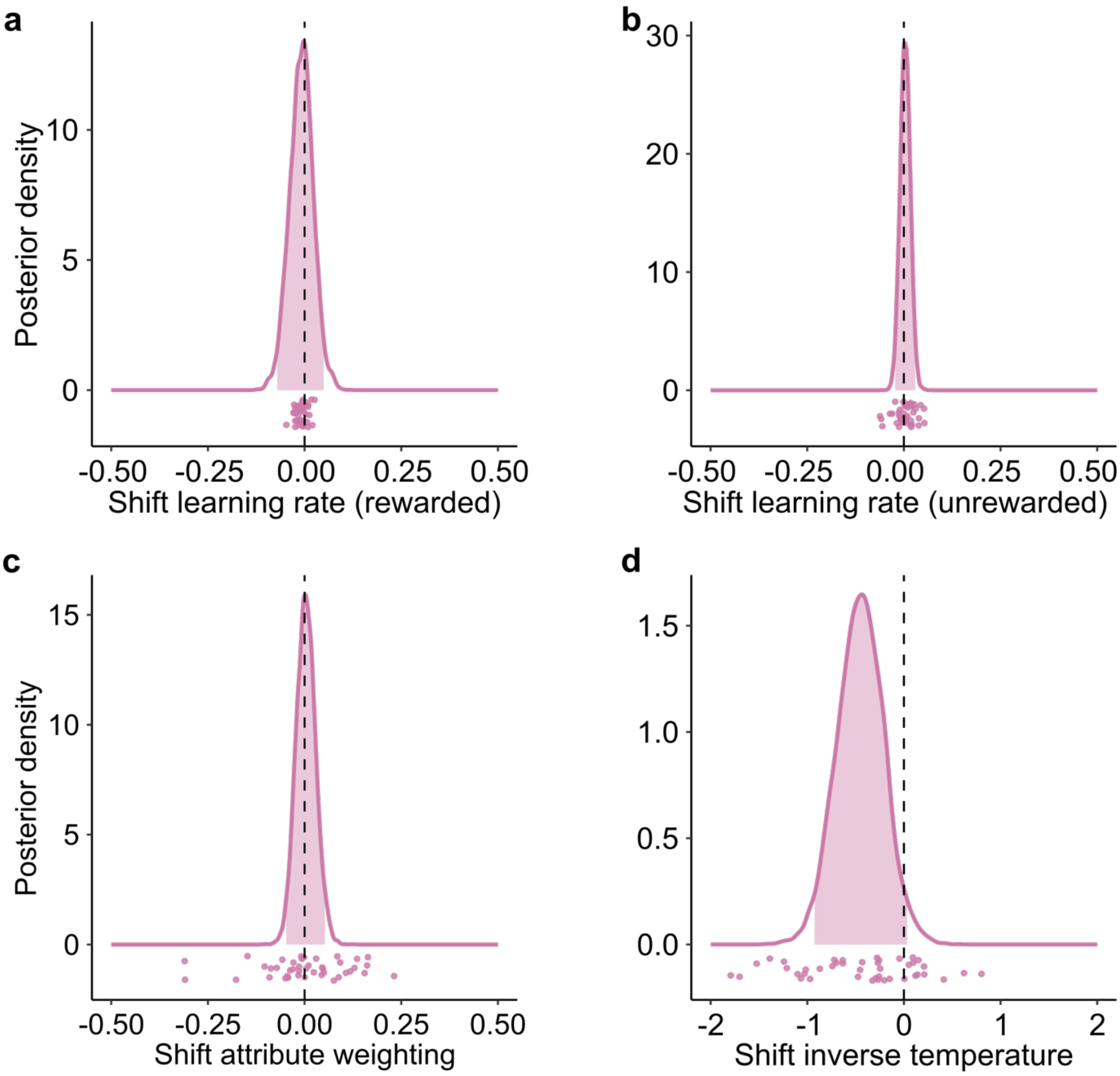

**Figure S7. Biperiden-induced shifts in the stable phase of the learning task from the Bayesian hierarchical model.** Density of posterior predictive distributions of the biperiden-specific shift in **a**, **b** learning rate in rewarded $s_{\lambda r}$ and unrewarded trials $s_{\lambda u}$, **c** attribute weighting $s_{\omega P}$, and **d** inverse temperature $s_{\zeta}$. Positive shifts represent an increase under biperiden relative to placebo. Shaded areas represent the 95 %-HDI of the posterior predictive distribution and points single-subject means.

**Table S11. Group-level parameter estimates of the stable phase of the learning task.** Median (Mdn), standard deviation (SD), and lower and upper bounds of the 95%-HDI interval are given. The model consisted of estimates for the learning rate of rewarded choices $\lambda_r$, the learning rate of unrewarded choices $\lambda_u$, the attribute weighting $\omega_P$, the inverse temperature $\zeta$, and the corresponding biperiden-specific shifts on these parameters $s_{\lambda r}$, $s_{\lambda u}$, $s_{\omega P}$, and $s_\zeta$.

| **Parameter** | **Mdn** | **SD** | **2.5 %** | **97.5 %** |
|---|---|---|---|---|
| $\lambda_r$ | *0.60* | *0.04* | *0.52* | *0.68* |
| $\lambda_u$ | *0.25* | *0.02* | *0.21* | *0.30* |
| $\omega_P$ | *0.59* | *0.04* | *0.50* | *0.66* |
| $\zeta$ | *5.58* | *0.43* | *4.80* | *6.47* |
| $s_{\lambda r}$ | -0.01 | 0.03 | -0.07 | 0.04 |
| $s_{\lambda u}$ | 0.00 | 0.01 | -0.02 | 0.03 |
| $s_{\omega P}$ | 0.00 | 0.03 | -0.05 | 0.05 |
| $s_\zeta$ | -0.45 | 0.24 | -0.92 | 0.03 |

### 6. MEG: ROI selection

Average power in rewarded versus unrewarded trials during (0 – 500 ms) and after (500 – 1000 ms) feedback presentation (figure S8).

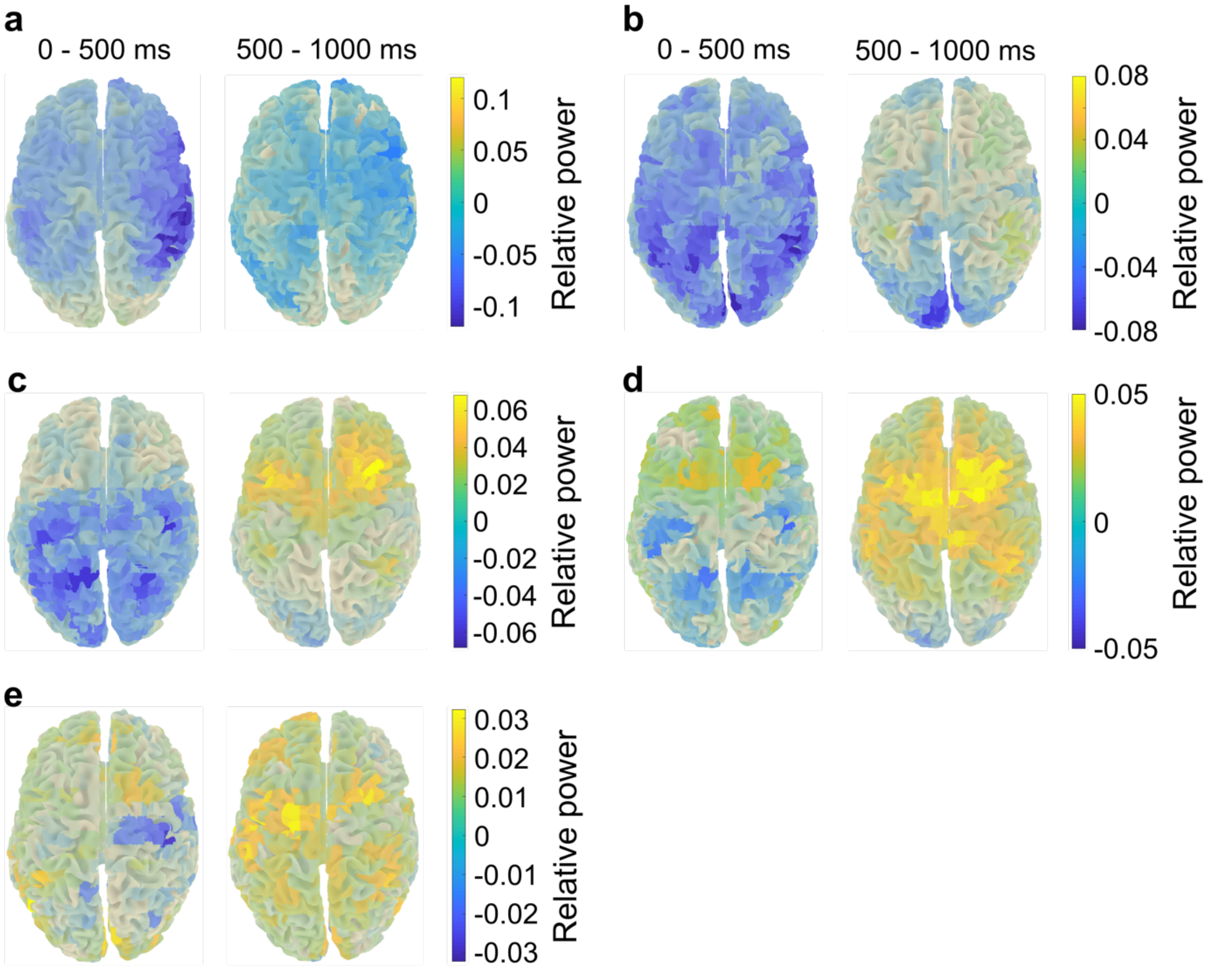

**Figure S8. Average power in rewarded versus unrewarded trials during (0 – 500 ms) and after (500 – 1000 ms) feedback presentation.** The power is averaged within the theta (4 – 8 Hz, **a**), alpha (8 – 13 Hz, **b**), low-beta (13 – 20 Hz, **c**), high-beta (20 – 30 Hz, **d**), and low-gamma band (30 – 40 Hz, **e**). Yellow represents higher power in rewarded relative to unrewarded trials, while blue represents lower power in rewarded relative to unrewarded trials.

### 7. MEG: Mixed-effects regressions

Full results of the mixed-effects regression for the ROI in the right hemisphere (Table S12), and left hemisphere (Table S14), and the post hoc analysis (Table S13).

**Table S12: Results from the mixed-effects regression with high-beta band power within the ROI in the right hemisphere as dependent variable.**

| | ***β*** | **SE** | ***df*** | ***t*** | ***p*** |
|---|---|---|---|---|---|
| *Intercept* | *$3.67 \cdot 10^{-2}$* | *$4.36 \cdot 10^{-3}$* | *182.3* | *8.41* | *< .001* |
| *Drug* | *$-1.12 \cdot 10^{-2}$* | *$5.01 \cdot 10^{-3}$* | *603.3* | *-2.23* | *.026* |
| *Estimated probability* | *$-1.12 \cdot 10^{-2}$* | *$4.58 \cdot 10^{-3}$* | *12270* | *-2.45* | *.014* |
| *Outcome* | *$3.70 \cdot 10^{-3}$* | *$1.19 \cdot 10^{-3}$* | *13760* | *3.10* | *< .002* |
| *Choice side* | *$9.49 \cdot 10^{-3}$* | *$1.17 \cdot 10^{-3}$* | *13770* | *8.09* | *< .001* |
| Previous choice side | $-4.39 \cdot 10^{-4}$ | $1.17 \cdot 10^{-3}$ | 13770 | -0.37 | .709 |
| *Drug x Estimated probability* | *$1.37 \cdot 10^{-2}$* | *$6.28 \cdot 10^{-3}$* | *11410* | *2.19* | *.029* |
| Drug x Outcome | $2.56 \cdot 10^{-4}$ | $1.70 \cdot 10^{-3}$ | 13760 | 0.15 | .880 |
| Drug x Choice side | $-2.67 \cdot 10^{-4}$ | $1.67 \cdot 10^{-3}$ | 13780 | -0.16 | .873 |
| Drug x Previous choice side | $-1.06 \cdot 10^{-4}$ | $1.67 \cdot 10^{-3}$ | 13780 | -0.06 | .950 |

**Table S13: Mixed-effects regression for the biperiden condition (post hoc analysis).**

| | ***β*** | **SE** | ***df*** | ***t*** | ***p*** |
|---|---|---|---|---|---|
| *Intercept* | *$2.54 \cdot 10^{-2}$* | *$4.60 \cdot 10^{-3}$* | *118.7* | *5.52* | *< .001* |
| Estimated probability | $2.72 \cdot 10^{-3}$ | $4.36 \cdot 10^{-3}$ | 6797 | 0.62 | .533 |
| *Outcome* | *$3.90 \cdot 10^{-3}$* | *$1.21 \cdot 10^{-3}$* | *6797* | *3.22* | *.001* |
| *Choice side* | *$9.18 \cdot 10^{-3}$* | *$1.20 \cdot 10^{-3}$* | *6777* | *7.69* | *< .001* |
| Previous choice side | $-6.08 \cdot 10^{-4}$ | $1.20 \cdot 10^{-3}$ | 6779 | -0.51 | .611 |

**Table S14: Results from the mixed-effects regression with high-beta band power within the ROI in the left hemisphere as dependent variable.**

| | ***β*** | **SE** | ***df*** | ***t*** | ***p*** |
|---|---|---|---|---|---|
| *Intercept* | *$2.78 \cdot 10^{-2}$* | *$4.12 \cdot 10^{-3}$* | *201.4* | *6.73* | *< .001* |
| Drug | $-6.43 \cdot 10^{-3}$ | $5.06 \cdot 10^{-3}$ | 386.4 | -1.27 | .205 |
| Estimated probability | $-3.67 \cdot 10^{-3}$ | $4.43 \cdot 10^{-3}$ | 13010 | -0.83 | .408 |
| *Outcome* | *$2.31 \cdot 10^{-3}$* | *$1.15 \cdot 10^{-3}$* | *13750* | *2.01* | *.045* |
| *Choice side* | *$-5.57 \cdot 10^{-3}$* | *$1.13 \cdot 10^{-3}$* | *13760* | *-4.92* | *< .001* |
| *Previous choice side* | *$2.72 \cdot 10^{-3}$* | *$1.13 \cdot 10^{-3}$* | *13760* | *2.40* | *.016* |
| Drug x Estimated probability | $8.28 \cdot 10^{-3}$ | $6.09 \cdot 10^{-3}$ | 12610 | 1.36 | .174 |
| Drug x Outcome | $-8.32 \cdot 10^{-4}$ | $1.64 \cdot 10^{-3}$ | 13760 | -0.51 | .612 |
| Drug x Choice side | $-3.04 \cdot 10^{-4}$ | $1.62 \cdot 10^{-3}$ | 13780 | -0.19 | .851 |
| Drug x Previous choice side | $-2.36 \cdot 10^{-3}$ | $1.62 \cdot 10^{-3}$ | 13780 | -1.46 | .143 |

### 8. Exclusion criteria

In our study, healthy male participants conducted behavioural tasks after an oral dose of a placebo and the cholinergic antagonist biperiden. Concurrently, MEG was recorded and after the MEG sessions, an anatomical MRI was acquired. To ensure both MEG/MRI compatibility and a good health status of the participants, we had a preceding screening session. Participants with indications found in List S1 were excluded from the study.

**List S1. Exclusion criteria.**

- Weight < 60 kg or > 90 kg
- BMI < 18 or > 28
- Systolic blood pressure > 140 mmHg
- Diastolic blood pressure > 90 mmHg
- BDI score > 12
- Impaired vision (and no contact lenses)
- Lactose intolerance
- Regular/recent use of drugs (incl. alcohol, cigarettes)
- Psychiatric diseases
- Neurological diseases
- Hepatic dysfunction
- Renal dysfunction
- Diseases of the cardiovascular system
- Epilepsy
- Glaucoma
- Thyrotoxicosis
- Gastrointestinal diseases
- Diabetes
- Metal implants
- Claustrophobia

### 9. Bayesian hierarchical models: posterior predictive checks

To assess whether the Bayesian hierarchical models could capture participants' behaviour, we conducted posterior predictive checks for the choice task and both phases of the learning task. Therefore, we simulated 500 datasets based on subject-level estimates of the parameters per participant. Then, we correlated the probability of high EV choice, high magnitude choice, and high probability choice for simulated and raw data.

#### 9.1. Gambling task

For the gambling task, the model could capture participants' behaviour (Figure S10a-c). Additionally, simulated behaviour and participants' behaviour were highly significant (Figure S10d-f, Table S15).

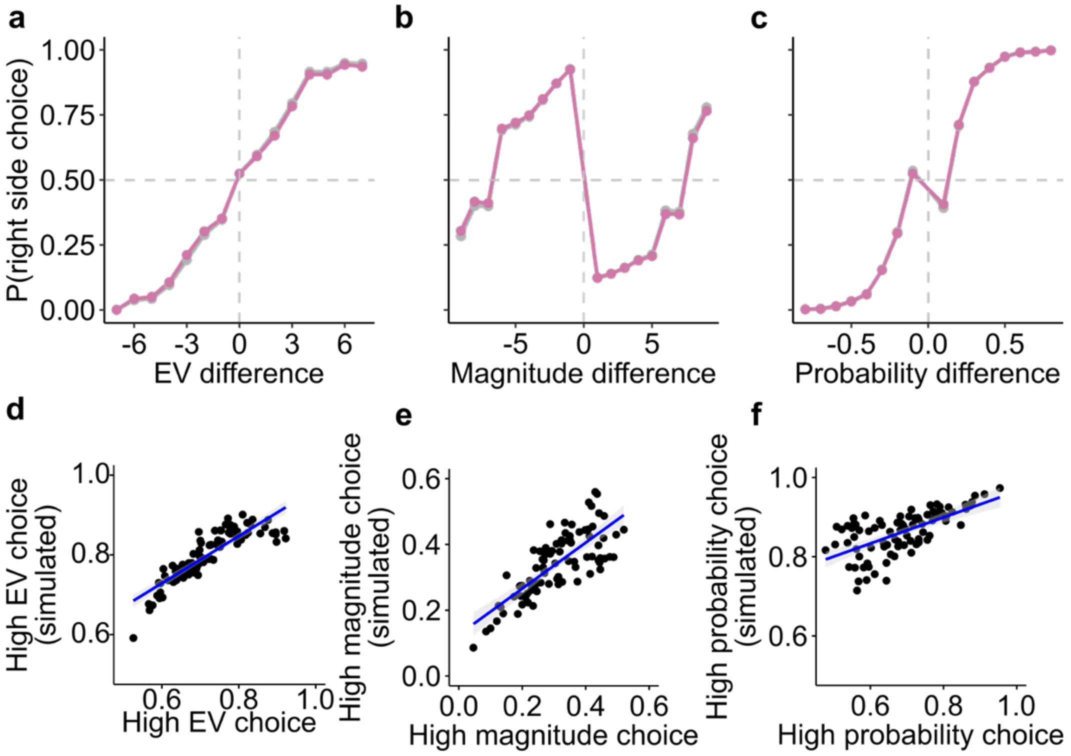

**Figure S9: Posterior predictive checks for the gambling task.** Probability of simulated right-side choice as a function of difference in **a** EV, **b** reward magnitude, and **c** reward probability between the two options. Choice behaviour was simulated for the biperiden session (pink) and placebo session (grey). Solid lines represent mean, shaded areas SEM across simulations. Correlation between participants' and simulated choices for **d** high EV option, **e** high magnitude option, and **f** high probability option.

**Table S15: Posterior predictive checks for the gambling task.** Correlation coefficients of participants' choices and simulated choices for EV, reward magnitude and estimated reward probability.

| **Task parameter** | ***R*** | ***p*** |
|---|---|---|
| *EV* | *0.9* | *< .001* |
| *Magnitude* | *0.76* | *< .001* |
| *Probability* | *0.66* | *< .001* |

### 9.2. Learning task: stable phase

The model for the stable phase of the learning task could capture participants' behaviour (Figure S10a-c). Additionally, simulated behaviour and participants' behaviour were highly significant (Figure S10d-f, Table S16).

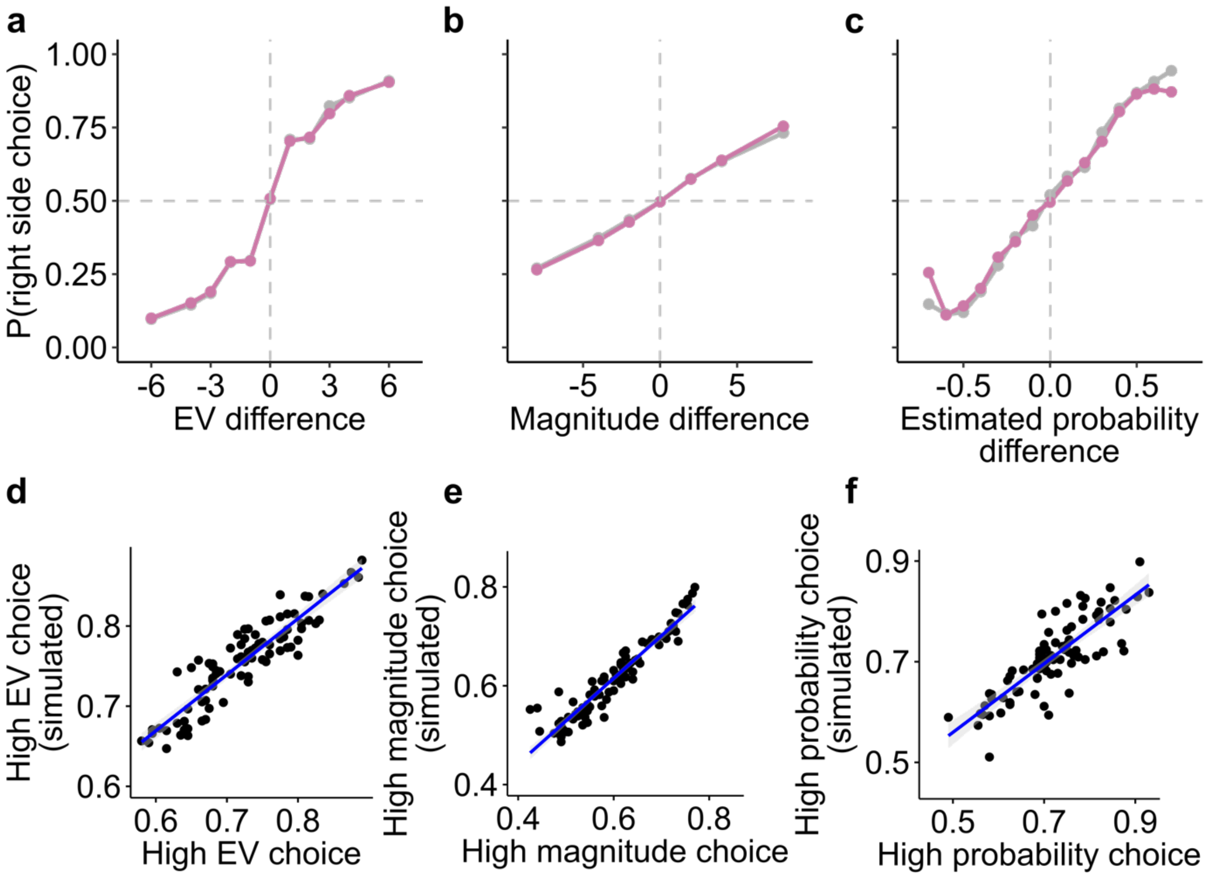

**Figure S10: Posterior predictive checks for the stable phase of the learning task.** Probability of simulated right-side choice as a function of difference in **a** EV, **b** reward magnitude, and **c** estimated reward probability between the two options. Choice behaviour was simulated for the biperiden session (pink) and placebo session (grey). Solid lines represent mean, shaded areas SEM across simulations. Correlation between participants' and simulated choices for **d** high EV option, **e** high magnitude option, and **f** high probability option.

**Table S16: Posterior predictive checks for the stable phase of the learning task.** Correlation coefficients of participants' choices and simulated choices for EV, reward magnitude and estimated reward probability.

| **Task parameter** | ***R*** | ***p*** |
|---|---|---|
| *EV* | *0.91* | *< .001* |
| *Magnitude* | *0.94* | *< .001* |
| *Probability* | *0.83* | *< .001* |

### 9.3. Learning task: volatile phase

The model for the volatile phase of the learning task could capture participants' behaviour (Figure S11a-c). Additionally, simulated behaviour and participants' behaviour were highly significant (Figure S11d-f, Table S17).

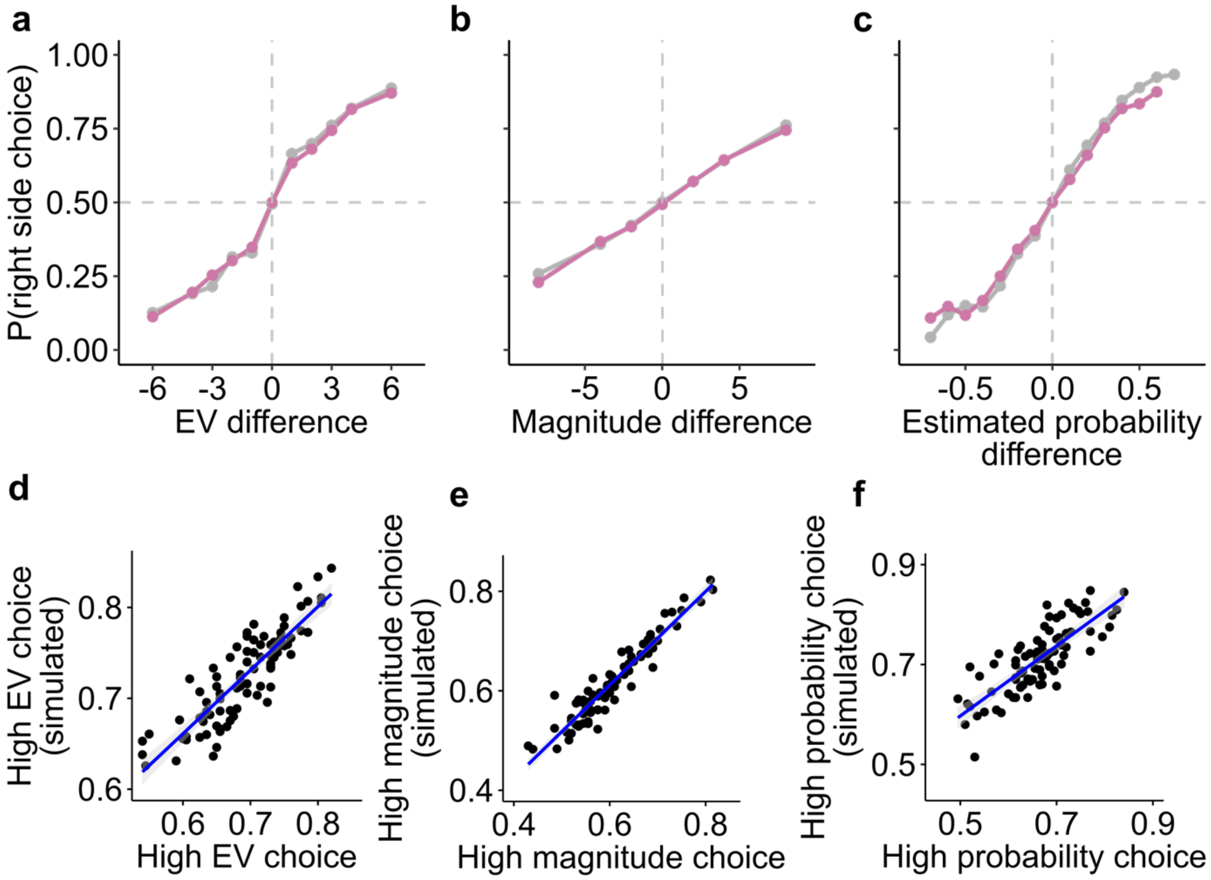

**Figure S11: Posterior predictive checks for the volatile phase of the learning task.** Probability of simulated right-side choice as a function of difference in **a** EV, **b** reward magnitude, and **c** estimated reward probability between the two options. Choices based on estimated reward probabilities were decreased under biperiden (pink) compared to placebo (grey). Solid lines represent mean, shaded areas SEM across simulations. Correlation between participants' and simulated choices for **d** high EV option, **e** high magnitude option, and **f** high probability option.

**Table S17: Posterior predictive checks for the volatile phase of the learning task.** Correlation coefficients of participants' choices and simulated choices for EV, reward magnitude and estimated reward probability.

| **Task parameter** | ***R*** | ***p*** |
|---|---|---|
| *EV* | *0.87* | *< .001* |
| *Magnitude* | *0.93* | *< .001* |
| *Probability* | *0.80* | *< .001* |

### 10. MEG: Time-resolved illustrations of drug effects

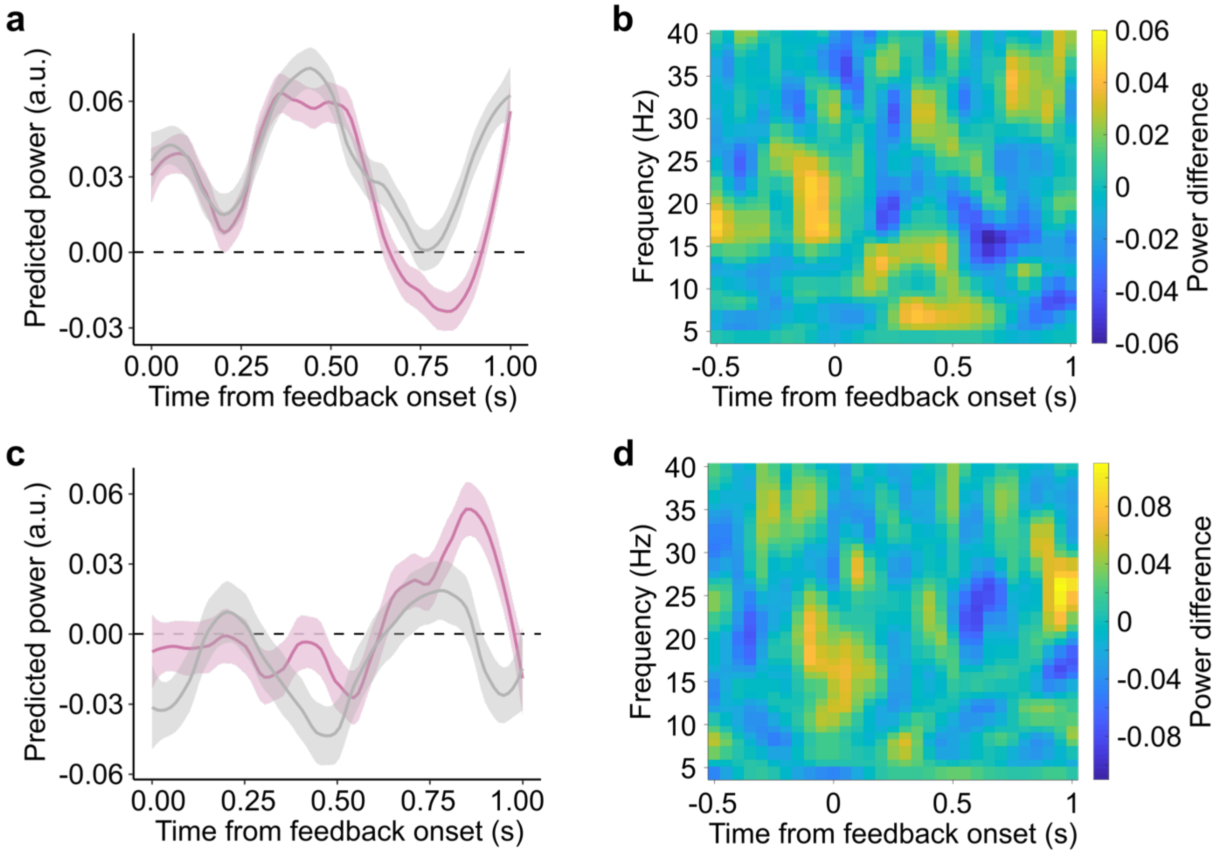

**Figure S12. Time-resolved illustrations of predicted high-beta power and time-frequency representations (TFRs) for the main effect of drug (a, b) and drug-by-estimated-probability interaction (c, d) averaged within the ROI. a** Predicted high-beta power as a function of time and drug as extracted from multiple mixed-effects regressions (step size of 0.05 s). **b** Model-independent TFR for biperiden minus placebo sessions averaged within the ROI (with lower power under biperiden relative to placebo marked in blue and higher power under biperiden relative to placebo marked in yellow). **c** Predicted high-beta power scaling with estimated probability as a function of time and drug as extracted from multiple mixed-effects regressions (drug x SP), step size of 0.05 s). **d** TFR of drug-dependent SP-modulation (colour convention as in b). Within the ROI, the power difference between trials with estimated probabilities in the upper and lower quartiles was calculated, and the difference of this contrast between the biperiden and placebo sessions is shown.
